# Developmental effects of oxytocin neurons on social affiliation and processing of social information

**DOI:** 10.1101/2020.10.08.330993

**Authors:** Ana Rita Nunes, Michael Gliksberg, Susana A.M. Varela, Magda Teles, Einav Wircer, Janna Blechman, Giovanni Petri, Gil Levkowitz, Rui F. Oliveira

## Abstract

Hormones regulate behavior either through activational effects that facilitate the acute expression of specific behaviors or through organizational effects that shape the development of the nervous system thereby altering adult behavior. Much research has implicated the neuropeptide oxytocin (OXT) in acute modulation of various aspects of social behaviors across vertebrate species, and OXT signaling is associated with the developmental social deficits observed in autism spectrum disorders, however, little is known about the role of OXT in the neurodevelopment of the social brain. We show that perturbation of OXT neurons during early zebrafish development led to a loss of dopaminergic neurons, associated with visual processing and reward, and blunted the neuronal response to social stimuli in the adult brain. Ultimately, adult fish whose OXT neurons were ablated in early life, displayed altered functional connectivity within social decision-making brain nuclei both in naïve state and in response to social stimulus and became less social. We propose that OXT neurons have an organizational role, namely to shape forebrain neuroarchitecture during development and to acquire an affiliative response towards conspecifics.

**Significance Statement:** Social behavior is developed over the lifetime of an organism and the neuropeptide oxytocin modulates social behaviors across vertebrate species, and is associated with neuro-developmental social deficits such as autism. However, whether oxytocin plays a role in the developmental maturation of neural systems that are necessary for social behavior remains poorly explored. We show that proper behavioral and neural response to social stimuli depends on a developmental process orchestrated by oxytocin neurons. Animals whose oxytocin system is ablated in early life show blunted neuronal and behavioral responses to social stimuli as well as wide ranging disruptions in the functional connectivity of the Social Brain. We provide a window into the mechanisms underlying oxytocin-dependent developmental processes that implement adult sociality.

## INTRODUCTION

Social behavior describes any of a wide group of behaviors by an organism towards members of its species, termed conspecifics (Robinson et al., 2019). The systems underlying these behaviors are thought to be established during development whereas the behavioral manifestations themselves are not immediately put into effect, but become apparent only later in life.

The “organizational hypothesis” claims that hormones can shape the structure of the developing nervous system and as a result alter the adult animal behavior and was first suggested by Phoenix, et al. following their study of the developmental effect of testosterone exposure on subsequent sexual behavior (Phoenix et al., 1959). Organizational effects refer to long-term, irreversible impact of hormones on tissue differentiation that can either directly or indirectly influence physiology, metabolism and behavior. In the context of behavior, organizational signals that can be mediated by neuropeptides, are necessary during critical developmental windows in order to shape neural systems whose activity will only become relevant at a later stage in the lifetime of the animal. This organizational effect is contrasted with the acute or “activational” effect exerted by hormones, neurotransmitters and neuropeptides on behavior, physiology and metabolism.

The neuropeptide oxytocin (OXT) has a widely-studied activational role in several aspects of social behavior (for a review see (Donaldson and Young, 2008; Grinevich et al., 2016; Jurek and Neumann, 2018)), including social processing (Gamer et al., 2010; Grinevich and Stoop, 2018), attention (Bartz et al., 2011; Lukas et al., 2011; Shamay-Tsoory and Abu-Akel, 2016), reward (Insel and Shapiro, 1992; Bowen and Neumann, 2017), and pro-social behaviors (Lukas et al., 2011), in rodents and humans. Previously, we showed that this activational role of OXT is evolutionarily conserved in deep time, since OXT also modulates perception of visual social cues (Nunes et al., 2020) and social recognition (Ribeiro et al., 2020a, 2020b) in zebrafish.

OXT is thought to have organizational effects as well. As early as 1989, Noonan et al. showed that pharmacological treatment of neonatal rats with OXT had long term effects on behavior in the adult (Noonan et al., 1989). In a more recent example, Keebaugh and Young utilized selective viral overexpression to modulate OXTR levels in the nucleus accumbens of pre-pubertal female prairie voles and showed that this developmental intervention is necessary and sufficient to induce alloparental behavior in the adult (Keebaugh and Young, 2011). Such results have been described in various mammalian models, behavioral domains, and using pharmacological (e.g. peptide, agonist and antagonist injection) as well as genetic interventions in prenatal and prepubertal animals (for a review see (Miller and Caldwell, 2015) and (Hammock, 2015)). Furthermore, human studies of Autism Spectrum Disorder (ASD) have strongly suggested that certain features of ASD arise from very early developmental abnormalities which are thought to be linked to OXT signaling, further suggesting an organizational role for OXT in the development of social behavior (Heinrichs et al., 2009; Hovey et al., 2014) or for a review (Guastella and Hickie, 2016). However, the mechanisms underlying this organizational mode of action have thus far remained elusive.

It has been well established that oxytocin communicates with other neurotransmitter systems for proper social behaviors. Dopamine (DA) is mainly involved in reward and reinforcement systems (Dölen and Malenka, 2014; Love, 2014), but also in sensory modulation and attention gating (Love, 2014; Grinevich et al., 2015). DA neurons are activated during social interactions and mating, and interact with the OXT system to promote pair-bond formation (Dölen and Malenka, 2014; Johnson et al., 2017) and sociability (Hung et al., 2017), suggesting that OXT and dopamine are both necessary to promote the sensory and rewarding aspects of social interactions.

Here, we show that in zebrafish, signaling by OXT neurons is required in an organizational manner during early life for the display of social affiliation in adulthood. Furthermore, we go on to show that perturbing this process in a very early developmental time window, equivalent to fetal developmental stages in mammals, has long-lasting effects on specific dopaminergic clusters associated with reward and visual attention/ processing, and also leads to wide-ranging changes in brain activity within previously described social processing networks.

## MATERIALS AND METHODS

### Experimental model

Zebrafish were raised and bred according to standard protocols. Zebrafish were kept in mixed sex groups (10 adults/l) in a recirculation life supporting system (tecniplast) with the following parameters: 28 °C, pH 7.0, conductivity 1000 µS/cm, 14 L:10D light:dark cycle. Fishes were fed with a combination of live food (Paramecium caudatum and Artemia salina) and commercial processed dry food (Gemma). Husbandry protocols, water chemistry and health program have been described previously (Borges et al., 2016). All experimental procedures were conducted in accordance with standard operating procedures of the Instituto Gulbenkian de Ciência and Direcção Geral de Alimentação e Veterinária (DGAV-Direcção Geral de Alimentação e Veterinária, permit number 0421/000/000/2015), Portugal, and Institutional Animal Care and Use Committee (IACUC, protocol number 27100516) of the Weizmann Institute, Israel.

Zebrafish transgenic/mutant lines used in this study: *Tg(oxt:EGFP*)(Blechman et al., 2011), *Tg(oxt:Gal4)wz06* (Anbalagan et al., 2018), *Tg(UAS:NTR-mCherry)c264* (Davison et al., 2007), *Tg(UAS:sypb-EGFP)* (Meyer and Smith, 2006).

Larvae and adult zebrafish (3-6 months old) of both sexes were used in this work.

### Social affiliation assay

The social preference test followed the protocol described previously (Wircer et al., 2017; Ribeiro et al., 2020a). Briefly, focal zebrafish were given a choice between two side-by-side compartments: one containing a shoal (two males and two females) and an empty one (Figure 1A-B) during a 10-min test. The stimulus shoal matched the genotype of the focal fish. The stimulus compartment was randomly switched between tests, to control for any place preference possibly induced by the arena or laboratory frames. All compartments were completely sealed to block transmission of chemical and vibrational stimuli and thus, only visual cues were accessible. The experimental test tank was placed over a custom-built infrared LED light box, to increase image quality for subsequent automated video tracking. Fish behavior was recorded from above with either a high-speed camera FLARE (2M360, Io Industries) or a B&W mini surveillance camera (Henelec 300B) connected to a computer, using video recording software (Pinnacle Studio 12). Videos were analyzed with Ethovision XT11.0 (Noldus Inc., The Netherlands). Relevant data were then exported and further analyzed. The two regions of interest, empty and shoal, were defined as the 1/10 of the length of the arena immediately adjacent to the empty compartment or the compartment containing the stimulus shoal, respectively. The percentage of cumulative time fish spent in these regions of interest (%T_shoal_ and %T_empty_) was used to calculate the social affiliation score [%T_shoal_/(%T_shoal_ +%T_empty_)]. This score, also called the sociability score, has been commonly used in rodent studies to measure sociability in the 3-chamber test (Park et al., 2018). Total distance traveled was also measured.

**Figure 1.**
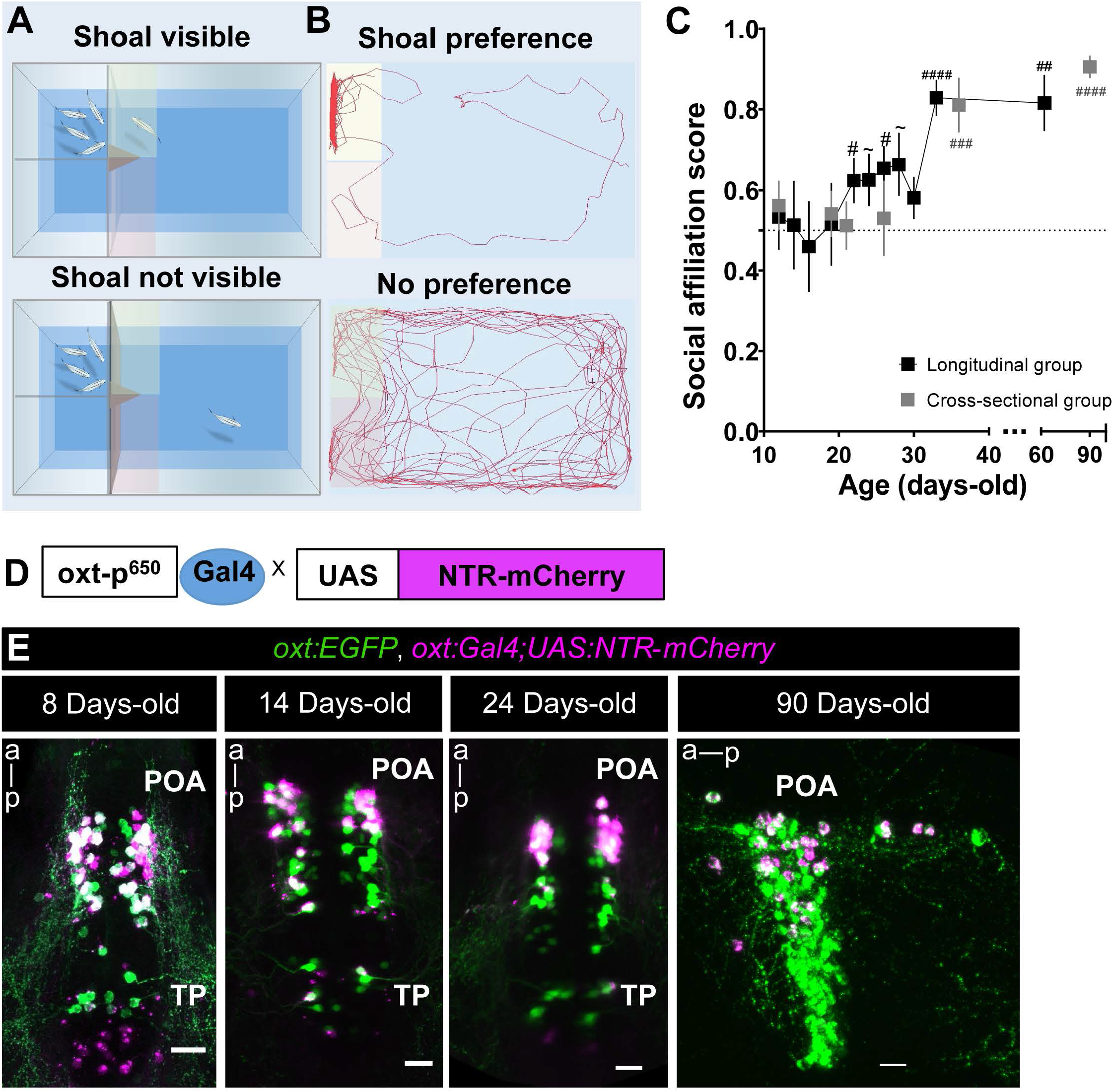
Behavioral and genetic tools to study the role of oxytocinergic neurons in adult social affiliation. (A,B) Schematic representation of the visual-mediated shoal preference behavioral test setup. (A) When a mixed-sex shoal stimulus is visible, focal fish spends most of the time near the shoal (top). When the shoal is not visible (bottom), the focal fish explores the entire arena equally. (B) Representative tracking of the focal fish behavior. (C) Ontogeny of social affiliation in zebrafish. Visual preference to associate with a shoal of conspecific fish (social affiliation score> 0.5) emerges after the third week of life. Social affiliation score = %T_shoal_/(%T_shoal_ + %T_non-shoal_). # Hash indicates a significant preference towards conspecifics, determined by a one-sample *t*-test with a hypothesized value of 0.5 (chance level). Fish were tested either repeatedly throughout development (black squares, longitudinal group) or only once at certain developmental time points (grey squares, cross-sectional group). (D) Spatio-temporal control of oxytocin-specific transgene expression. *oxt*:Gal4 drives the expression of nitroreductase tagged with mCherry fluorescent protein (NTR-mCherry) in oxytocinergic neurons. (E) *Tg(oxt:Gal4;UAS:NTR-mCherry)* characterization: NTR-mCherry+ cells co-localized with a subpopulation of *oxt*:eGFP neurons, at all developmental stages studied (8, 14 and 24 days-old: whole-mount larvae confocal z-stack image, dorsal view, anterior to top; 90 days-old, close-up of the preoptic area (POA), sagittal brain slice, confocal z-stack image, anterior to left). Scale: 20 μm. TP, Posterior Tuberculum; POA, Preoptic area. Data presented as mean ± SEM. # p<0.05, ## p<0.01, ### p<0.001, #### p<0.0001 (black # relates to the longitudinal group, grey # relates to the cross-sectional group).

#### Ontogeny of social affiliation behavior

Social preference was tested at different developmental stages: early larvae (12 days-old), mid larvae (14-20 days-old), post-flexion metalarvae (21-30 days-old), juveniles and adults (from 30 to 90 days-old). The size of the testing arena for early larvae was 3.5 x 6.0 cm; for mid larvae 7.0 x 8.0 cm; for metalarvae 7.0 x 10.0 cm; and for juveniles and adults it was 20.0 × 20.0 cm. The stimulus shoal matched the developmental stage of the focal fish. Two different experiments were performed: a) longitudinal, where individual fish were repeatedly tested at each developmental time point, and b) cross-sectional, where individual fish were tested only once, at a given developmental time point.

From all behavioral experiments, animals were excluded from analysis if they failed to enter both the ROIs (i.e. Shoal and Empty) as this results in a social score equals to infinity.

### Chemically induced OXT neuronal ablation

#### Early ablation

Nitroreductase (NTR)-mediated cellular chemically induced OXT neuronal ablation was conducted as previously described(Curado et al., 2007, 2008). Briefly, larvae (∼30-40 per petri dish) were treated with metronidazole (MTZ, Vetranal CAT#46461, Sigma) dissolved in Danieau buffer to a final concentration of 10mM for 48 h while being protected from light, to prevent MTZ photoinactivation. After the first 24 h, the MTZ-containing Danieau buffer was replaced by fresh MTZ medium. Control untreated larvae were subjected to the same procedure in Danieau buffer without MTZ, but were also protected from light. After 48 h of treatment, larvae were washed out several times in Danieau buffer.

#### Adult ablation

We used a modified ablation protocol of 3X48 h MTZ treatments (5 mM), as we observed that this increased survival of adult animals. Briefly, adult zebrafish were placed into 2-liter tanks containing MTZ (Veterinary preparation, Vetmarket; Shoham, Israel, CAT#165228, or 2-methyl-5-nitroimidazole-1-ethanol, TCL Europe, CAT#M0924) dissolved in system water at a final concentration of 5 mM, protected from light. After ∼16 h of treatment, zebrafish were moved to a tank containing fresh system water without MTZ, fed, and allowed to recover for ∼2 h, followed by a second treatment of MTZ 5mM for 16 h. Animals were then allowed to recover in their home tanks for 48 h. The entire protocol was repeated three times.

#### Effect of chemically induced OXT neuronal ablation on endogenous OXT cells

Zebrafish treated with MTZ at different time points during development (4-6 days-old, 12-14 days-old, 20-22 days-old or 90 days-old) were allowed to recover from treatment for 48 h and were then sacrificed to assess the treatment effects on endogenous OXT cells by *in situ* hybridization (see below) or by direct visualization of the mCherry transgene signal. OXT cells were manually counted using imageJ software (Schindelin et al., 2012), except for *in situ* experiment at 22-24 days-old where we used a semi-automated method for quantifying total fluorescence (see below).

#### Recovery of endogenous OXT cells following early-life chemically induced OXT neuronal ablation

Zebrafish larvae were ablated at 12-14 DPF, and then allowed to grow until adulthood. At 90 DPF animals were sacrificed, brains harvested, and subjected to fluorescent in situ hybridization (see below). The total fluorescence was then quantified in a semi-automated manner (see below).

#### Effect of chemically induced OXT neuronal ablation on adult social affiliation

Zebrafish treated with MTZ at 4-6 days-old, 12-14 days-old, 20-22 days-old or 90 days-old were allowed to grow until adulthood to be tested for social affiliation behavior (see above).

#### Time course of NTR-mCherry recovery following 4-6 days-old chemically induced OXT neural ablation

Fish were treated with MTZ at 4-6 days-old and sampled at different time points: 8, 12, 19, 26 and 42 days-old.

#### Semi-automated method for quantifying total fluorescence

For quantification of images in the recovery experiment, and the effect of ablation at 22-24 DPF on OXT mRNA we utilized a semi-automated method for quantifying total fluorescence. Each Z-stack was broken into its component optical planes, and ROI’s were selected for the region containing the cells and the background (a region within the tissue with no fluorescent cells). The threshold for “positive” pixels was defined as the 75th percentile value of al the pixels in the background ROI, and all positive pixels per slice were counted. the total number of positive pixels per sample was summed, and then normalized to the maximal value for that cohort. Source code in MATLAB for this procedure is included.

#### Effect of 4-6 days-old chemically induced OXT neuronal ablation on adult neuronal activity (pS6 activation)

To assess the effect of early OXT ablation on adult brain activation patterns in response to a social stimulus, adult zebrafish that had been treated with MTZ at 4-6 days-old and untreated controls were exposed individually to either a mixed-sex shoal of conspecifics or an empty tank for 10 min. Immediately after, we blocked the visual stimulus, without disturbing the focal fish, by placing an opaque partition between the experimental and stimulus tanks. After 50 min (to allow for expression of the pS6 neural activation marker), zebrafish were sacrificed and heads were collected and processed for paraffin slice immunofluorescence (see below).

#### Effect of 4-6 days-old chemically induced OXT neuronal ablation on larvae and adult dopaminergic system

Untreated and 4-6 days-old MTZ-treated zebrafish larvae were allowed to recover from treatment (48h) and were either processed immediately for whole-mount larvae TH immunofluorescence (see below) or allowed to grow until adulthood and then processed for paraffin slice TH-immunofluorescence (see below).

### In situ hybridization

RNA *in situ* hybridization was performed on both whole larvae and whole adult brains. Larvae or dissected brains were fixed in 4%PFA and *in situ* hybridization was performed as described in (Machluf and Levkowitz, 2011; Wircer et al., 2017). An OXT probe was generated using a pGEM plasmid encoding for *oxt* mRNA (RefSeq NM_178291.2). Following development with Fast Red reagent (Roche, cat. No. 11496549001), adult brains were embedded in agar and sagittally cut at a thickness of 150μm on a vibratome, and whole larvae and brain slices were mounted in glycerol and imaged on a Zeiss LSM 800 scanning confocal microscope.

### Whole-mount larvae immunofluorescence

Briefly, larvae were euthanized in ice-cold water, transferred to 4% PFA and then incubated overnight at 4°C on a shaker. Then, the PFA was washed out and samples were placed in pre-cooled acetone in a freezer at −20°C for 10 min. The acetone was washed out (PBS 0.1% TritonX-100), and the samples were then incubated in blocking solution (PBS 0.1% triton + 1% DMSO + 1% BSA + 5% NGS) for minimum of 2 h at room temperature (RT), followed by incubation with primary antibody overnight at 4°C on a shaker. Primary antibodies used were either mouse anti-TH (MAB 318, Merck Millipore), rabbit anti-EGFP (A11122, ThermoFisher) or guinea pig anti-OXT (T-5021, Peninsula labs), at a concentration of 1:200. Next, samples were washed repeatedly (minimum of 6X15 min washes) with blocking solution and then placed in a blocking solution containing fluorescent secondary antibody (1:200) overnight at 4°C on a shaker. Then, samples were washed in PBS, mounted dorsally on a slide in mounting medium (Aqua-Polymount, Polysciences Inc., Warrington PA, CAT# 18606-20) and imaged by a Zeiss LSM 800 scanning confocal microscope.

### Paraffin adult brain slice immunofluorescence

Zebrafish heads were fixed in 10% buffered formalin for 72 h and decalcified in EDTA (0.5 M, pH 8.0) for 48 h, followed by paraffin inclusion. Coronal slices (6 μm-thick) were cut with a microtome. Sectioned slices were then processed for immunofluorescence. After antigen retrieval with Tris-EDTA (10 mM Tris Base, 1 mM EDTA, 0.05% Tween20) at 95°C for 20 min, slices were washed in TBS-Tx (TBS 0.025% TritonX-100, 3X10 min), then incubated with blocking solution (TBS 0.025% TritonX-100 + 1%BSA) for 1 h at RT, followed by an overnight incubation with a primary antibody (1:400 at 4°C). Slides were then washed in TBS-0.025% TritonX-100 (3X10 min) and incubated in secondary antibody (1:1000 for 2 h). After washes, slices were incubated with DAPI for 20 min, then rinsed in TBS and mounted with EverBrite^TM^ Hardset Mounting medium (Biotium). Primary antibodies used were anti-phosphorylation of S6 ribosomal protein, pS6 (Cell signaling S235/236) and anti-tyrosine hydroxylase, anti-TH, (mouse monoclonal, MAB 318, Merck Millipore).

### Neuronal quantifications

#### Neuronal pS6 cell quantification

Coronal images were acquired with a commercial Nikon High Content Screening microscope, based on Nilon Ti equipped with a Andor Zyla 4.2 sCMOS camera, using a 20X 0.75 NA objective, quadruple dichroic filter, and controlled with the Nikon Elements software. To avoid double-imaging of the same cells, we imaged every other slice. We quantified density of pS6-positive cells in 16 brain areas that belong to the social decision making network (O’Connell and Hofmann, 2011): Vv-ventral nucleus of the ventral telencephalic area (V), homologous to the mammalian lateral septum, subdivided into: anterior (Vv_a) and posterior (Vv_p); Vd-dorsal nucleus of V, homologous to the mammalian Nacc, subdivided into: anterior (Vd_a), medial (Vd_m) and posterior (Vd_p); Vc-central nucleus of V, homologous to the mammalian striatum, subdivided into: anterior (Vc_a) and posterior (Vc_p); Dm- medial zone of the dorsal telencephalic area (D), homologous to the mammalian bLAMY; Dl-lateral zone of D, homologous to hippocampus; Dd-dorsal zone of D; Vs-supracommissural nucleus of V, homologous to the mammalian extended amygdala (BNST and meAMY); Vp-postcommissural nucleus of V; Ppa-parvocellular preoptic nucleus, anterior part and Ppp-parvocellular preoptic nucleus, posterior part, both homologous to the mammalian preoptic area; VM-ventromedial thalamic nucleus; VL-ventrolateral thalamic nucleus (Figure 6-1).

For each brain area, about five coronal brain slices were analyzed manually using imageJ software (Schindelin et al., 2012). Background subtraction and linear adjustments of brightness and contrast levels were performed in the same way for all groups. The brain areas of interest were identified using DAPI to define neuroanatomical boundaries and landmarks identified in the Zebrafish Atlas (Wullimann and Mueller, 2004). In each brain slice, we placed one square of 1000 μm^2^ in the brain region of interest, in each hemisphere, following these criteria: the square was always placed over the highest number of pS6-positive cells, keeping minimum distance from the border and edge of the brain section, a similar strategy that has been used by others (Lorenzi et al., 2017). pS6-positive cells were counted if a nucleus surrounded by the cytoplasm was clearly visible, and if the intensity of the pS6 signal was perceptibly greater than background (see Figure 6 for examples of pS6 staining in the sampled brain areas).

#### Dopaminergic neuronal quantification

Dopaminergic quantification was performed in both larvae and adult zebrafish, either 4-6 days-old MTZ-treated or untreated. Briefly, for larvae, whole-mount zebrafish immunofluorescence was performed as described above using primary antibody anti-TH. Larvae were imaged and confocal z-stacks were analyzed by ImageJ. Larva dopaminergic forebrain, pretectum and posterior tuberculum nucleus were identified according to (Sallinen et al., 2009; Tay et al., 2011) and TH+ cells were counted in these three groups.

For adults, heads from both treatment groups (untreated and 4-6 days-old MTZ-treated) were collected and processed for paraffin embedding, followed by paraffin slice immunofluorescence as described above. Coronal slices were acquired with a SlideScanner Zeiss AxioScan Z1 (Zeiss), and analyzed with Zeiss Zen Lite software. To avoid double-imaging of the same cells, we imaged every other slice. Boundaries separating brain nuclei and subdivisions were identified based on DAPI staining, using as reference a coronal atlas of the zebrafish (Wullimann et al., 1996). Dopaminergic neurons (TH^+^) were identified using the same criteria as for pS6^+^ neurons.

TH^+^ nuclei groups were identified according to (Rink and Wullimann, 2001; Panula et al., 2010; Parker et al., 2013) (Figure 4-1). We counted TH^+^ cells in the following clusters: the subpallium area (extending from the rostroventral telencephalon to the dorsocaudal telencephalon, also identified as G2 by Panula and colleagues (Panula et al., 2010), which included a TH cluster containing Vv, Vc and Vl, and in-between areas, a TH cluster including Vd and the lateral area outside the Vd, and a TH cluster within the Vs (identified by the anterior commissure) and lateral area outside the Vs; TH clusters in the PPa area were divided in two: a more anterior (close to the border of the diencephalon, lateral margin of the ventricle, G3) and a more posterior extending area in the medial area of the PPa (ventral part of the PPa), G4); a TH cluster in the pretectum (PPr, G7); in the ventromedial and ventrolateral nucleus (Vm, Vl, G6); a TH cluster in the periventricular nucleus of the posterior tuberculum (TP/TP, small cells, ventral to the central posterior thalamic nucleus (CP), G11) and in the posterior tuberal nucleus (PTN, G12) (Figure 4-1). Similar to pS6 neuronal quantification described above, we manually analyzed five coronal brain slices for each cluster, in each brain hemisphere, placing one square of 1000 μm^2^ in the brain region of interest with the highest density of cells.

### Statistical analysis

Data are represented as mean ± standard error of the mean (SEM). Normality of the data was tested by D’Agostino and Pearson omnibus normality test and Shapiro-Wilk normality tests. When parametric assumptions were verified, we used parametric statistics; otherwise we used equivalent non-parametric tests, namely Mann-Whitney tests to compare the effect of MTZ treatment on OXT mRNA, on the expression of the NTR-mCherry transgene, and to compare the adult social affiliation score between untreated and MTZ-treated groups at different developmental time points (4-6 days-old, 12-14 days-old, 22-24 days-old and 90 days-old MTZ-treatment). Significance was denoted as p < 0.05, and p-values refer to two-tailed tests, unless otherwise noted. The use of one-tailed tests was justified by a priori directionality hypotheses in the following cases:

- Effect of MTZ treatment on OXT mRNA and NTR-mCherry transgene expression (Figure 2 and Figure 2-1), since in our system, we expected that MTZ would ablate the NTR-oxytocinergic expressing neurons;
- Effect of MTZ treatment (oxytocinergic ablation) on social affiliation behavior in fish expressing the NTR transgene, and thus causing OXT ablation, since OXT is well known to regulate social behaviors and thus, we expected that oxytocinergic ablation would lead to a decreased social behavior in zebrafish (Figure 2).

**Figure 2.**
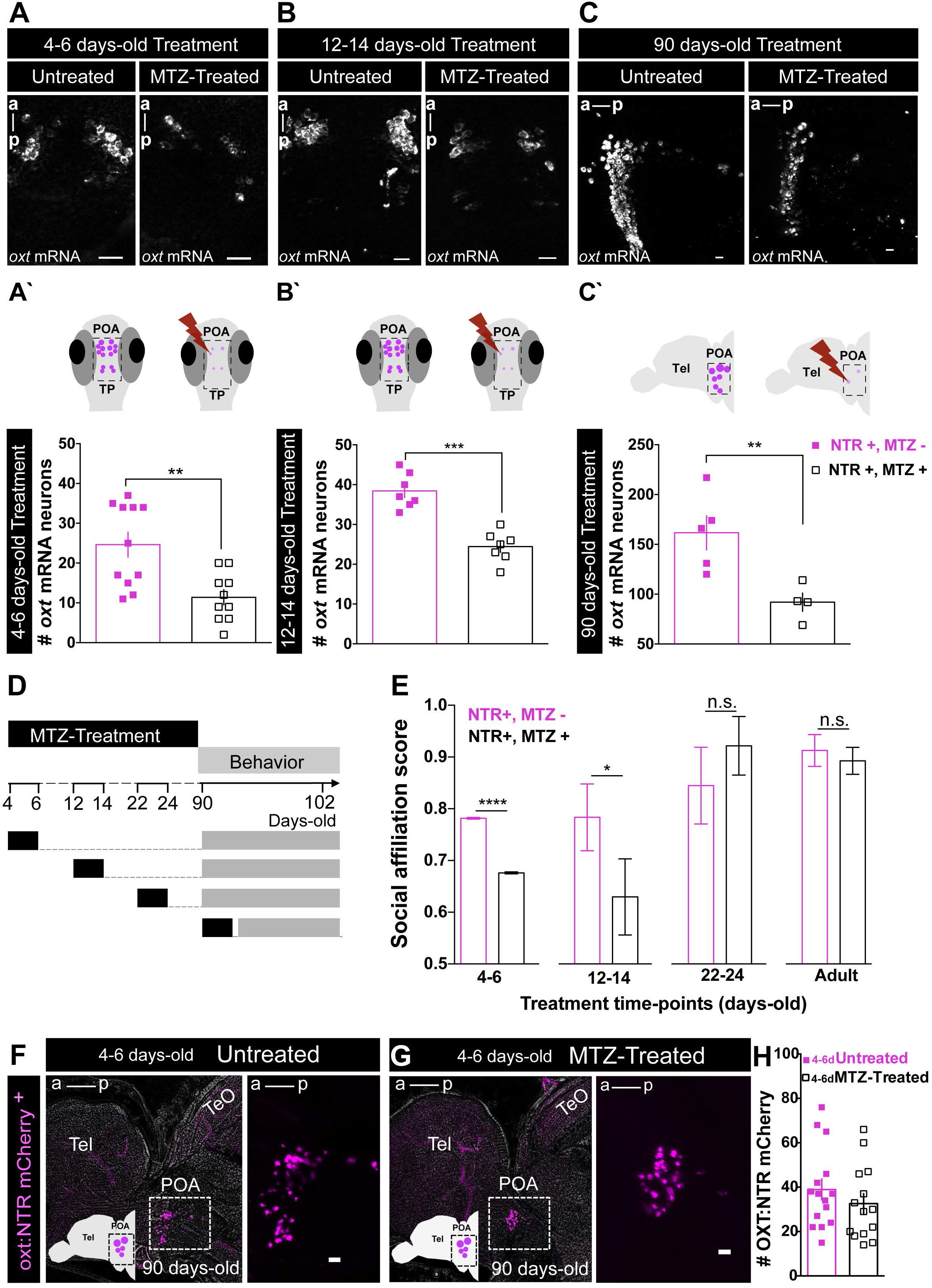
Contrasting organizational vs activational effects of oxytocin neurons in adult social affiliation. (A-C) Representative examples of the effects of metronidazole (MTZ)-induced ablation on endogenous OXT as detected by *in situ* hybridization, at different treatment time-points. (A) 4-6 days-old treatment and (B) 12-14 days-old treatment: whole-mount larvae confocal maximum intensity z-stack image, dorsal view, anterior to top; (C) 90 days-old treatment: sagittal brain slice, confocal maximum intensity z-stack image, anterior to left. (A’-C’) Quantification of the number of cells expressing OXT mRNA in untreated fish or following MTZ treatment at: (A’) 4-6 days-old treatment, (B’) 12-14 days-old treatment and (C’) 90 days-old treatment. (D) Experimental timeline: fish were MTZ-treated at different developmental stages, allowed to grow until adulthood and then tested for social affiliation. (E) MTZ treatments during the first two weeks of life affect adult social affiliation. Social affiliation score = %T_shoal_ / (%T_shoal_ + %T_non-shoal_). Eight independent cohorts were treated at 4-6 days-old, and only one cohort at later time points. Therefore, in order to obtain comparable sample sizes among the different age-treatment groups and a more representative sample, the data for 4-6 days-old MTZ treatment shown in this graph were generated by a Monte Carlo simulation with 1000 iterations of samples of 15 individuals of each group (4-6 days-old MTZ-treated vs. untreated; see Statistical analysis subsection for details). One-tailed p-values were used because there was an *a priori* directional hypothesis that MTZ-treatment would ablate OXT neurons in fish expressing the NTR transgene and induce a decrease in social affiliation behavior. (F-H) OXT neurons ablated at 4-6 days-old were recovered by adulthood. (F) Representative example of NTR-mCherry-expressing OXT neurons in untreated adult fish. (G) Representative example of NTR-mCherry-expressing OXT neurons in adult fish treated with MTZ at 4-6 days-old. Images in (F,G) are maximum intensity confocal z-stacks, sagittal slices, anterior to left. (H) Quantification of the number of NTR-mCherry cells in untreated (full squares) and 4-6 days-old MTZ-treated adult fish (empty squares). Scale: 20 μm. Data presented as mean ± SEM. Full squares: untreated fish (NTR+, MTZ); Open squares: MTZ-treated fish (NTR+, MTZ+). *p<0.05, **p<0.01,*** p<0.001. ****p<0.0001.

#### Statistical analysis of behavioral data

One sample *t*-tests were used to verify if social affiliation during development was statistically different from chance (0.5). Because we tested eight independent cohorts for the effect of MTZ-treatment at 4-6 days-old time point (by different researchers in two different laboratories), the total sample size for this time point (n=210) was much larger than for the other time points (12-14 days-old: n=28, 22-24 days-old, n=19; adult: n=13). Taking in account all eight cohorts together, the adult social affiliation score of the 4-6 days-old MTZ-treated fish were significantly decreased from untreated siblings (p=0.0012, Mann-Whitney test, n= 112 untreated vs. 98 MTZ-treated fish). However, a Generalized Linear Model (GLM) with beta regression and planned comparisons, which was used to compare the effect of MTZ treatment on adult social affiliation score of the eight different cohorts (Figure 2-3), revealed not only an effect of the MTZ treatment (F_1_=9.486, p=0.0021) but also variation among cohorts (F_7_=2.719, p=0.0081, Figure 2-4). We therefore performed a Monte Carlo simulation for a sample size of 30 individuals sampled from the observed data, 15 individuals from each group (untreated and 4-6 days-old MTZ treated) with 1000 iterations. This simulation, ensured a more comparable sample size for the 4-6 days-old time point treatment relative to other time-point treatments. Moreover, it ensured a more representative sample than each cohort independently or the total eight-cohort population. We obtained a Monte Carlo p-value of p<0.001 (Figure 2E), which means that there was a significantly higher number of simulations in which the social affiliation score of the untreated group was greater than the social affiliation score of the MTZ-treated group, as compared to the number of simulations with the inverse trend, for a sample size of 15 individuals, across all eight cohorts. P-value was calculated as the mean social affiliation score of the 1000 iterations for untreated fish divided by the mean scores of the 1000 iterations for treated fish divided by 1000.

To determine whether the difference in social affiliation score between untreated and 4-6 days-old MTZ-treated fish was due to OXT ablation and not movement impairments caused by MTZ, we used a Linear Model (LM) to compare the total distance moved (square root transformed) of both groups in the eight independent cohorts. Both GLM with beta regression on adult social affiliation score and LM on total distance moved (square root transformed) were also applied to different control cohort fish not expressing NTR transgene (Figure 2-3, 2-5). In all of these models (GLM and LM) the explanatory variables were treatment and cohorts for mains effects and interaction.

#### Statistical Analysis of neuronal activation (ps6 cell quantification)

To compare neuronal activation between treatments and stimuli, we used a Generalized Linear Mixed Model (GLMM) with a Poisson distribution using the brain identification number as the random effect in the statistical model, followed by planned comparisons, since we target areas that are known to belong to the social decision-making network. We checked whether the data were normally distributed with Shapiro-Wilk tests for each brain area. Since they were not, and because the data are counts, we checked the fit of the data to the Poisson distribution with the “fitdistrplus” package from R (Delignette-Muller and Dutang, 2015). By visual inspection of density and cumulative distribution plots we validated that the Poisson distribution was the most appropriate to our data analyses. As fixed effects we used treatment and stimuli. We analyzed the 16 brain areas independently and corrected the p-values of the planned comparison with the false discovery rate (FDR) adjustment method.

#### Statistical Analysis of dopaminergic cell quantification

To compare dopaminergic cell numbers between treatments, and since the data were normally distributed (G7: W = 0.97685, p-value = 0.6383; G12: W = 0.9546, p-value = 0.1459; G2: W = 0.95265, p-value = 0.1267; Shapiro-Wilk test), LM was performed to assess the effect of 4-6 days-old MTZ-treatment in total TH counts of three distinct independently analyzed TH clusters in 8 days-old larvae (Figure 3, 3-1).

**Figure 3.**
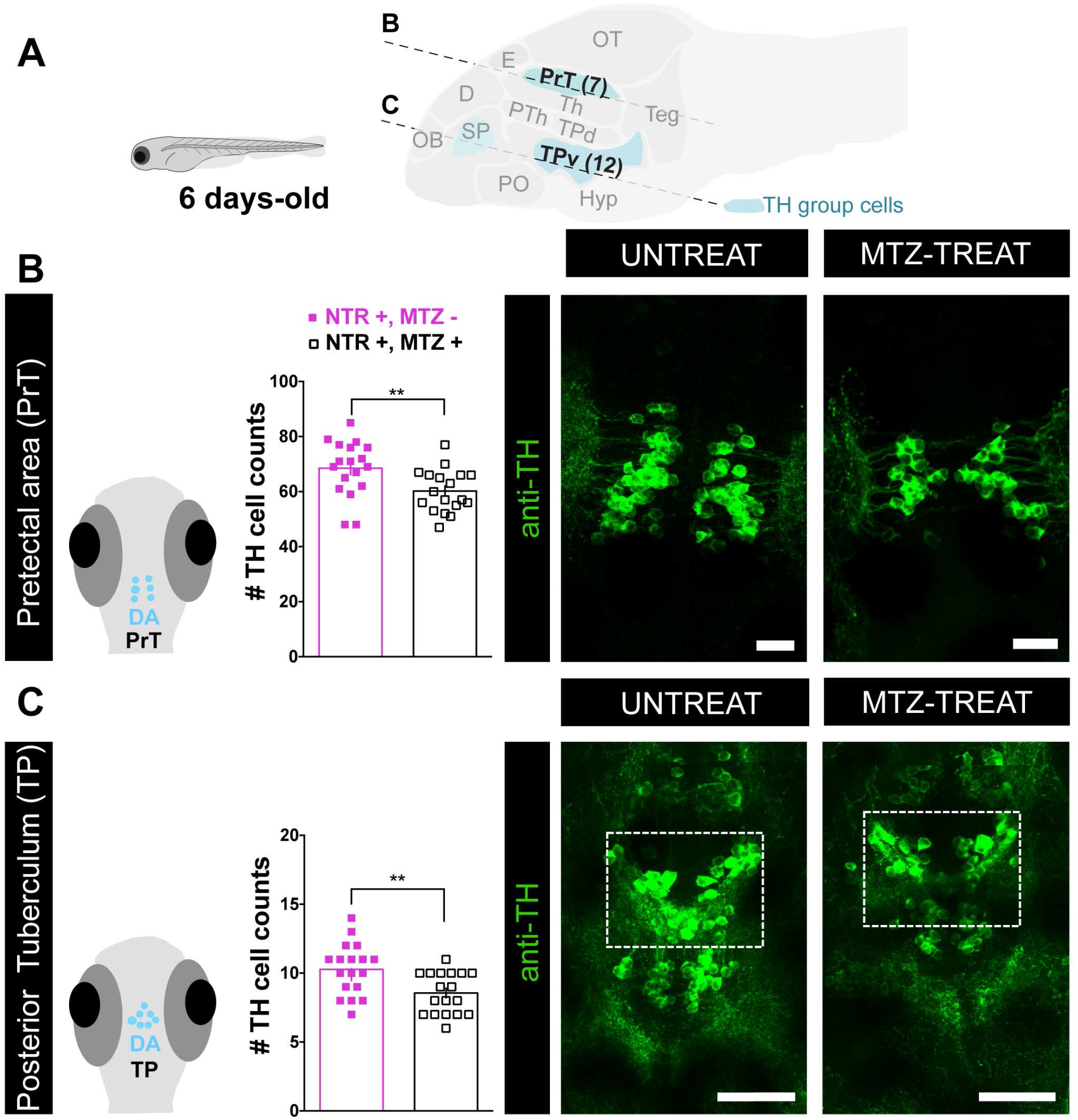
Early-life OXT ablation affects the dopaminergic system in early development. (A) Top: Schematic sagittal section through a zebrafish larva brain showing the three dopaminergic clusters that were analyzed in this study, and highlighting two key dopaminergic brain areas that were affected by OXT neuronal ablation at 4-6 days-old, namely the pretectum (PrT) and posterior tuberculum (TP). Schematic was adapted from *zebrafishucl.org/forebrain-regions/posterior-tuberculum*. (B) Quantification of total pretectum (PrT) TH cell counts in untreated (NTR+, MTZ-) vs 4-6 days-old MTZ-treated (NTR+, MTZ+) larvae and respective representative images. Scale: 20 μm. (C) Quantification of total Posterior Tuberculum (TP) TH cell counts in untreated (NTR+, MTZ-) vs 4-6 days-old MTZ-treated (NTR+, MTZ+) larvae and respective representative images. Scale: 50 μm. Data presented as mean ± SEM. Full squares: untreated fish (NTR+, MTZ); Open squares: MTZ-treated fish (NTR+, MTZ+). *p<0.05, **p<0.01,*** p<0.001. ****p<0.0001. *D*, dorsal telencephalon; *E*, epiphysis; *OB*, olfactory bulb; *OT*, optic tectum; *PO*, preoptic area; *PrT*, pretectum; *TPd*, posterior tuberculum dorsal part; *PTh*, prethalamus; *TPv*, posterior tuberculum ventral part; *Teg*, tegmentum; *Th*, thalamus; *SP*, subpallium (includes Vv, Vd and Vs in adult); *Hyp*, hypothalamus; *NTR*, nitroreductase; *MTZ*, metronidazole.

The effect of 4-6 days-old MTZ treatment on adult dopaminergic system was also assessed with a GLMM with a Poisson regression followed by planned comparisons, comparing the number of TH-positive cells sampled in five squares of 1000 µm^2^ per TH cluster in 8 different adult brain TH clusters. As fixed effects we used treatment and stimuli. All eight areas were analyzed independently and p-values of the planned comparisons were corrected with the FDR adjustment method (Figure 4 and 4-2).

**Figure 4.**
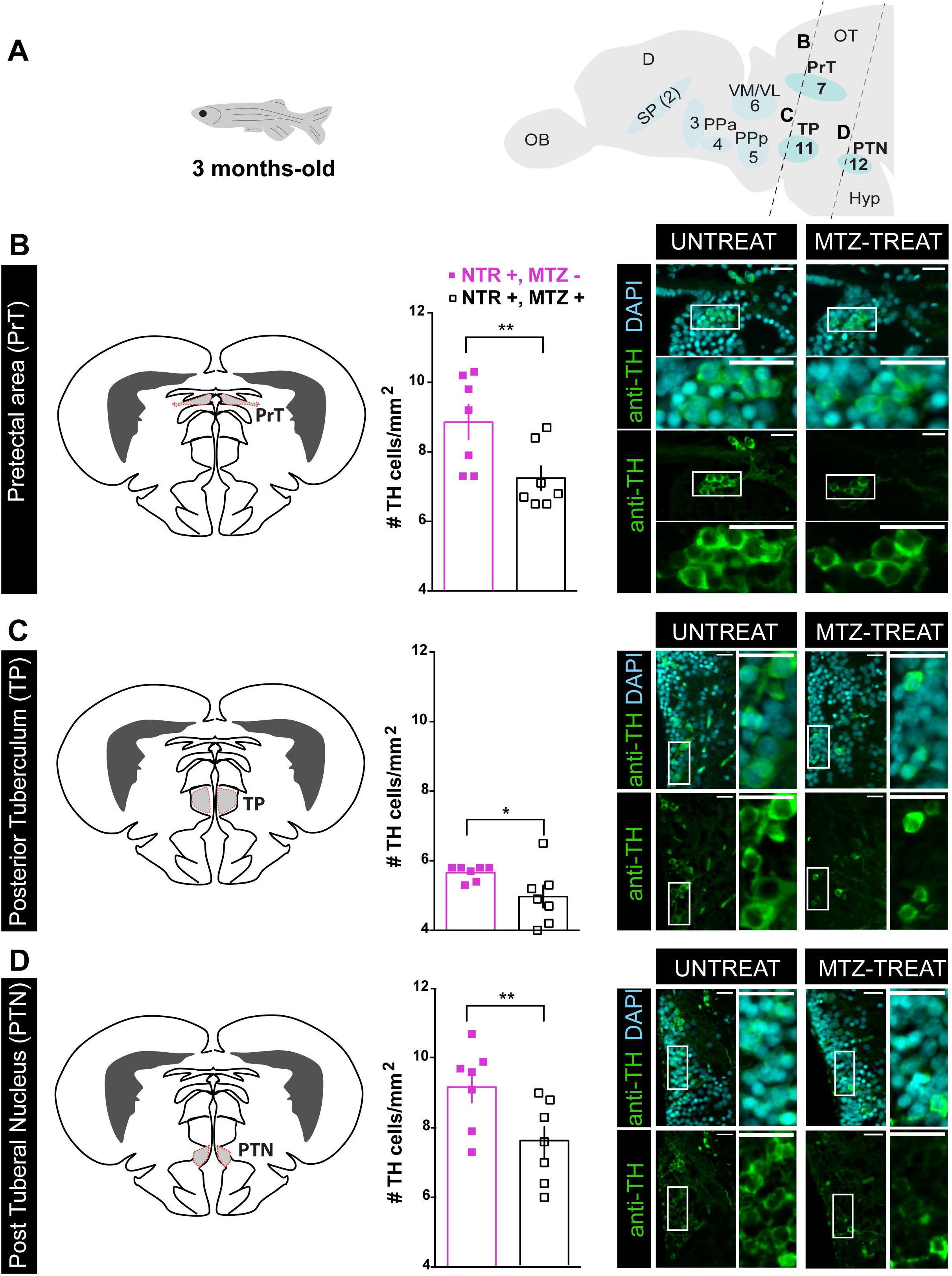
Early-life OXT ablation affects the dopaminergic system in adulthood. (A) Schematic sagittal section through a zebrafish adult brain, showing the eight dopaminergic clusters analyzed in this study, and highlighting the areas that were affected by OXT neuronal ablation at 4-6 days-old: Pretectal area (PrT), Posterior Tuberculum (TP) and Posterior Tuberal Nucleus (PTN). Dopaminergic groups are named according to (Panula et al., 2010; Parker et al., 2013). (B-D) Quantification of TH cell density (cells/mm^2^; mean of five sampled slices for each area, see Methods section for details) in brains of adult fish untreated (NTR+, MTZ-) or MTZ-treated at 4-6 days-old (NTR+, MTZ+) and respective representative images in: (B) Pretectal area (PrT), (C) Posterior Tuberculum (TP) and (D) Posterior Tuberal Nucleus (PTN) Scale: 20 μm. Data presented as mean ± SEM. Full squares: untreated fish (NTR+, MTZ); Open squares: MTZ-treated fish (NTR+, MTZ+). *p<0.05, **p<0.01,*** p<0.001. ****p<0.0001. *D*, dorsal telencephalon; *OB*, olfactory bulb; *OT*, optic tectum; *PrT*, pretectum; *TP*, posterior tuberculum; *SP*, subpallium (includes Vv, Vd and Vs); *Ppa*, anterior parvocellular preoptic nucleus; *VM*, ventromedial thalamic nucleus; *Vl*, ventrolateral thalamic nucleus; *PPp*, posterior parvocellular preoptic nucleus; *PTN*, posterior tuberal nucleus; *Hyp*, hypothalamus; *NTR*, nitroreductase; *MTZ*, metronidazole.

#### Data analysis software

Graphical representations of the data were performed in GraphPad Prism 6.0c software. Shapiro-Wilk normality test, one-sample *t*-test and *t*-tests were performed in GraphPad Prism, and the remaining tests were performed in R programming software, version 3.6.3 (R_core_Team, 2013) with the following packages: Ime4(Bates et al., 2015) and afex (Henrik Singmann, Ben Bolker, Jake Westfall, 2020) (for GLMM with Poisson regression), betareg (for GLMM with beta regression), emmeans (for planned comparisons)(Lenth, 2020), and the base R package (for LM regression and Monte Carlo simulations)(R_core_Team, 2013).

### Functional connectivity analysis

Functional connectivity, which has been defined as the temporal coincidence of spatially distant neurophysiological events (Friston, 1994), is typically inferred from temporal correlations between distinct brain areas in human fMRI data from both resting state and task-state studies (Biswal et al., 1997). The rationale for this approach is that areas that show a consistent correlation in their activity are components of the same brain network. Therefore, the concept of functional connectivity is purely correlative in nature, and per se it does not imply a direct causal relationship or the occurrence of structural connectivity between the correlated brain areas (Eickhoff and Müller, 2015). Functional connectivity can also be inferred from correlations across subjects using the expression of molecular markers such as immediate early genes (Teles et al., 2015). Similarly, we computed functional connectivity for our neuronal activity data based on the co-expression of pS6 across different brain regions for the set of individuals of each experimental treatment. For this purpose we have developed the method described below, which statistically validates the identified networks.

#### Network reconstruction

To test for functional connectivity of early MTZ-treated vs untreated brains exposed to social stimuli, we reconstructed Pearson correlation matrices using a resampling procedure, similar to the Quadratic Assignment Procedure (QAP) (Makagon et al., 2012; Borgatti et al., 2013). Instead of defining a single network, we construct the set of possible networks obtained leaving out some of the specimens’ information. More precisely, consider the case of *M* specimens, each with an associated expression vector *x_i_* ∈ ℝ*^N^*, where *N* is the number of brain regions. For a given *m* < *M*, we consider all the 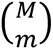 combinations Ωγ of *m* specimens and compute the corresponding functional graphs. We refer to the collection of graphs obtained in this way as a graph tower Ω_Γ_, where each of the combinations can be considered as a graph layer. The advantage of this construction is that each layer in the graph tower represents a different instance of the network bootstrapping. In this way, observables computable on a single layer can be bootstrapped across multiple ones. The construction has naturally one parameter, the sampling number *m*, which needs to be chosen on the basis of data-driven considerations. In our experiments, robustness analysis shows that results are robust for *m* values between 10 and 15. To obtained sparse functional connectivity matrices, we threshold each instance following (De Vico Fallani et al., 2017) at a density threshold of ρ = 0.2, which also corresponds to the minimum network heterogeneity across instances. After thresholding, for each treatment we average the thresholded instances to obtain a single matrix per treatment, which we use in the following analysis.

#### Detection of robust functional modules

Communities were computed using the Leiden community detection method (Traag et al., 2019) on the averaged treatment matrices. To increase the robustness of the detection, for each condition, we repeat the community detection 400 times. From the 400 candidate partitions we extract the central partition as described in (Peixoto, 2021) and associate the resulting central partition to the treatment under analysis. To quantitatively characterize differences among partitions, we measure the ratio *r* of total edge weights within a community with that of the edges between communities. More specifically, for a partition 𝒫 with *s* communities we compute the *s* × *s* matrix *P*, defined as

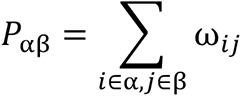

Where α, β = 0, … *m* − 1 label the modules of 𝒫 and ω*_ij_* is the edge weight between regions *i a*nd *j*. We then compute the ratio of average intra-community to inter-community edge weights as follows:

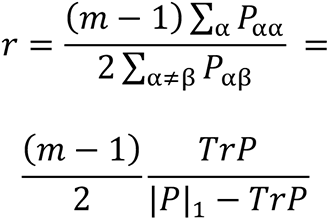

which measures the ratio of the average weight on the diagonal of *P*_αβ_ to the average diagonal weight. To assess significance of *r* values and of differences between them, we employ a permutation test based on permutating the community labels while preserving the size of the considered communities. We find that all *r* values are significantly different from zero, and so are also the differences in *r* between treatments (p<0.01).

#### Strength centrality

As a measure of local integration, we computed also the strength centralities. The first encodes how strongly a node links to its neighbors. while the second total measures how influential a node is at the network level on the basis of its direct connections and how well connected its neighbors are. In Figure 6-4 we report the nodes ranked in decreasing order of strength (weighted degree) centrality.

## RESULTS

### Early life ablation of oxytocinergic neurons decreases social affiliation

Zebrafish is a highly gregarious species exhibiting well-characterized social behaviors (reviewed in (Nunes et al., 2017)). We quantified the visually-mediated motivation of zebrafish to approach conspecifics, as an indicator of social affiliation, by performing a two-choice preference test measuring the time that individuals spend either near a compartment containing a shoal of conspecifics or an empty one (Ribeiro et al., 2020a, 2020b). When an adult zebrafish is visually presented with both shoal and non-shoal compartments, in a side-by-side configuration, it will spend most of the time in association with the shoal; however, if shoal is not visible, fish tend to explore the entire arena (Figure1 A,B). In accordance with previous reports (Engeszer et al., 2007; Dreosti et al., 2015), we found social affiliation is an acquired developmental trait that emerges after the third week of life. This was observed both in zebrafish that were repeatedly exposed to the social affiliation behavioral arena throughout development (12 days-old (d): p= 0.69, n=15; 14d: p=0.91, n=14; 16d: p=0.73, n=15; 16d: p=0.88, n=17; 22d: p=0.049, n=12; 24d: p=0.07, n=18; 26d: p=0.01, n=17; 28d: p=0.057, n=13; 30d: p=0.17, n=8; 33d: p<0.0001, n=14; 62d: p=0.002, n=10; one sample t-test vs. theoretical score of 0.5 indicating no preference, Figure 1C), and in zebrafish that were only exposed to the arena once at a specific developmental time point to avoid habituation to the setup (12d: p=0.34, n=16; 19d: p=0.48, n=18; 21d: p=0.85, n=6; 26d: p=0.76, n=10; 36d: p=0.0007, n=13; 90d: p<0.0001, n=24; one sample t-test vs. theoretical score of 0.5 indicating no preference, Figure 1C).

Oxytocin (OXT) has long been known to regulate social behaviors across species (Goodson, 2008) and it has been linked to neurodevelopmental disorders that impact social behavior during development and adulthood (Pobbe et al., 2012; Hovey et al., 2014). This makes OXT a good candidate system for investigating neurodevelopmental processes linked to social development. To disentangle the organizational vs. activational effects of OXT neurons in the acquisition of adult social affiliation, we used a transgenic line, *Tg(oxt:Gal4;UAS:NTR-mCherry)*, to express nitroreductase (NTR) protein fused to the mCherry reporter, in oxytocin neurons (Figure 1D,E). In the presence of the drug metronidazole (MTZ), NTR produces cytotoxic metabolites, thereby allowing temporally controlled ablation of OXT neurons at different developmental stages (Curado et al., 2007, 2008). We confirmed the expression of the NTR-mcherry fusion protein in the OXT-ergic neuronal population by co-localizing the *Tg(oxt:Gal4;UAS:NTR-mCherry)* with a transgenic reporter *Tg(oxt:EGFP)* at different stages of development (8, 14, 24 and 90 days-old, Figure 1E).

In both mammals and fish, OXT is produced by specific cells in the preoptic hypothalamus which can be classified as magnocellular or parvocellular, based on soma size (Sawchenko and Swanson, 1982; Van den Dungen et al., 1982; Knobloch and Grinevich, 2014; Grinevich et al., 2016). We and others have previously shown that zebrafish magnocellular OXT cells are clustered in the anterior-dorsal portion of the cell group, whereas the parvocellular neurons make up the more ventral part of the group (Wircer et al., 2017; Wee et al., 2019). We observed that the NTR-mCherry protein was prominently expressed in the most anterior-dorsal part of the preoptic area (Figure 1E), thus it can be considered to localize mainly to magnocellular OXT neurons in both larvae and adult zebrafish. These cells have been shown to project to ventral forebrain in fish (Saito et al., 2004). Also, recent research in rats has shown that these magnocellular cells project to several brains regions including hypothalamus, amygdala, lateral septum and nucleus accumbens (Zhang et al., 2021), indicating that this projection pattern is evolutionarily conserved. MTZ treatment at all time points was highly effective (>80% decrease) at ablating NTR-mCherry-positive OXT neurons (Figure 2-1). To ensure proper ablation of endogenous OXT-ergic cells following MTZ treatment, we also performed fluorescent *in situ* hybridization with probes directed against *oxt* mRNA at 4-6 days-old, 12-14 days-old, 22-24 days-old and in the adult (Figure 2A-C and Figure 2-2). A significant decrease in OXT-expressing cells was observed following MTZ treatment at all tested ages (p= 0.0024 n=11/10, p=0.0003 n=7/7, p=0.0011 n=7/8, p=0.0079 n=5/4, for 4-6 days-old, 12-14 days-old, 22-24 days-old and adult ablation untreated/treated, respectively, one tailed Mann-Whitney test, Figure 2A’-C’ and Figure 2-2).

To assess the effects of temporal OXT neuronal ablation on adult social affiliation, zebrafish were treated with MTZ at specific time-points during development (4-6, 12-14, 22-24 days-old) or post-puberty (90 days-old) and social affiliation was assessed in adulthood in comparison to untreated siblings (Figure 2D). We analyzed the social affiliation scores of eight independent cohorts in which OXT neurons were ablated at 4-6 days days-old and found that despite considerable variation among cohorts during the test, there was a robust effect of MTZ treatment on adult social affiliation (cohorts: p= 0.0081; treatment: p=0.001, see Statistical analysis section for detailed statistics of the Generalized Linear Model (GLM) with beta regression, Figure 2-3, Figure 2-4). MTZ treatment did not affect overall fish locomotion as measured by the total distance traveled during the trial (cohorts: p=8.342e-16, treatment: p=0.668, Linear Model (LM), Figure 2-3, Figure 2-4). We also observed variation among control adult cohorts not expressing NTR but treated or untreated with MTZ at 4-6 days-old, without a main effect of MTZ-treatment on either social affiliation score (cohort: p=0.0001, MTZ-treatment: p=0.2653, GLM with beta regression, Figure 2-3, Figure 2-5) or total distance moved (cohort: p<2e-16, MTZ-treatment: p=0.1536, LM, Figure 2-3, Figure 2-5).

In view of the biological variability observed across cohorts, we further verified the effect of early-life OXT-neuron ablation on social affiliation by utilizing a Monte Carlo (MC) simulation where we randomly sampled 15 treated and 15 untreated 4-6 days-old larvae from all 8 cohorts and repeated the procedure for 1000 iterations. This analysis also clearly indicated an impairment of adult social affiliation (p<0.0001, one-tailed Mann-Whitney test, Figure 2E, see Statistical analysis for details on MC), which was maintained in 12-14 days-old MTZ-treated adult fish (p=0.034, n=14/14, one-tailed Mann-Whitney test, Figure 2E).

In contrast to the deficit in social affiliation caused by early life MTZ treatment, ablation of OXT neurons in either 22-24 days-old or 90+ days-old animals did not affect adult social affiliation (22-24: p=0.1364, n=10/9 one-tailed Mann-Whitney test; Adults: p=0.3566, n=6/7, one-tailed Mann-Whitney test, Figure 2E). This result coincides with the observation that social affiliation is a developmentally acquired trait which is established before the juvenile stage of development (Figure 1C and (Dreosti et al., 2015).

Taken together, these results show that early developmental ablation of OXT neurons leads to a long-term deficit in social affiliation in adulthood, suggesting that OXT neurons are involved in an early-life developmental process that is required for proper social affiliation later in life.

### Early life OXT ablation leads to impairments in specific dopaminergic clusters

Notably, although early-life OXT neuronal ablation induced an impairment in social affiliation, we observed complete recovery of the OXT neural population by adulthood, as ablated fish displayed normal counts of OXT-expressing NTR-mCherry transgene (4-6 days-old MTZ-treated (n=14) vs. untreated (n=15) adult zebrafish: p= 0.33, unpaired t-test, Figure 2F-H). We verified this result, obtained from imaging of the NTR-mCherry transgene, by fluorescent in situ to OXT mRNA in adult fish subjected to early-life ablation, and found no significant difference from controls (MTZ-treated (n=4) vs. untreated (n=5) adult zebrafish: p= 0.25, two-tailed Mann-Whitney U test, Figure 2-7). We performed a time-course analysis for the OXT neural recovery and we found that already within 1 month (i.e. by 42 days-old) from the MTZ-treatment there is no discernible difference between treated and control animals (8 days-old, p<0.0001, n=15/15; 12 days-old, p<0.0001, n=13/7; 19 days-old, p<0.0001, n=10/9; 26 days-old, p=0.0048, n=12/11; 42 days-old, p=0.59, n=9/10; Unpaired t test with Welch’s correction, Figure 2-6). This is in line with previous work showing that zebrafish neurons are capable of regenerating following lesions even in adulthood (Kizil et al., 2012). Conversely, animals in which OXT neurons were ablated in adulthood displayed normal social drive (Figure 2E) despite the fact that the ablated cell populations had not yet recovered by the time of the behavioral testing (Figures 2C, 2-1).

We therefore hypothesized that the developmental organization of the dopaminergic system that also regulates social affiliation, might have been affected by early OXT ablation. Therefore, we examined whether early perturbation of the OXT neuronal system affected specific tyrosine hydroxylase (TH)-positive DA-ergic neuronal clusters (Figures 3, 4 and 4-1), which in zebrafish are anatomically discernible from other TH-positive catecholaminergic cells (Ma, 1994a, 1994b, 1997; Rink and Wullimann, 2001, 2002), and whether these effects persist over the long term. We observed that already 24 hours after ablation of OXT neurons at 4-6 days-old, there was a significant decrease in dopaminergic neuronal counts in the pretectum (PrT; p=0.009, n=18/18, LM, Figure 3B, Figure 3-1), and the large neurons of the posterior tuberculum (TP; p=0.0045, n=18/18, LM, Figure 3C, Figure 3-1). In contrast, no differences were found in telencephalic DA neurons of the ventral subpallium area (Figure 3-1). The pretectum is the teleostean homologue of the superior colliculus, an area known to be involved in gaze control and possibly attention in zebrafish (Antinucci et al., 2019) as well as in mammals (Krauzlis et al., 2004). The posterior tuberculum is the source of the ascending DA system, which is considered analogous to the mammalian ventral tegmental area (VTA) (Rink and Wullimann, 2001), which is involved in reward and reinforcement (Morales and Margolis, 2017).

Unlike the ablated OXT cells, which recovered over time, the deficits in DA neurons observed in early OXT-ablated larvae persisted through adulthood, as adult fish that had been treated with MTZ between 4-6 days-old displayed a reduced number of TH-positive DA cells in the PrT (p=0.005, n=7/7; Figure 4B, Figure 4-2, GLMM). Similarly, OXT ablation caused a decrease in the DA-ergic neurons residing in two subdivisions of the TP, the periventricular nucleus of the posterior tuberculum (TP) and the posterior tuberal nucleus (PTN), which are distinguishable only in the adult (TP p= 0.025, n=7/7; PTN p= 0.005, n=7/7; Figure 4C-D, Figure 4-2, GLMM with Poisson regression, planned comparisons and FDR corrections).

To demonstrate that at the time of their ablation, OXT neurons physically interact with the affected DA neurons, we examined whether OXT neurons form synapses on DA-ergic clusters of the PrT and TP (Figure 5). We employed a double transgenic line, *Tg(oxt:Gal4;UAS:Sypb-EGFP)*, which genetically expresses the synaptic marker, synaptophysin-GFP in OXT neurons (Anbalagan et al., 2019), and visualized synaptic contacts onto TH positive DA-ergic neurons (Figure 5). To ensure that these are indeed bona-fide OXT releasing synapses, we also co-stained these fish with an anti-OXT antibody (Blechman et al., 2018; Anbalagan et al., 2019) (Figure 5C-D). We found OXT projections forming multiple synapses onto the DA neurons of the PrT and the TP (Figure 5C,D1,D2). We also observed OXT cells directly abutting the TH-positive cells of the TP (Figure 5D3,D4), raising the possibility that OXT affects nearby DA-ergic cells in a non-synaptic manner as has been shown in other species (Ludwig and Leng, 2006; Son et al., 2013). These results indicate that these two systems, DA and OXT, are already linked at early developmental stages and that early ablation of OXT neurons irreversibly impairs the development of subsets of DA neurons which are key for social affiliation later in life.

**Figure 5.**
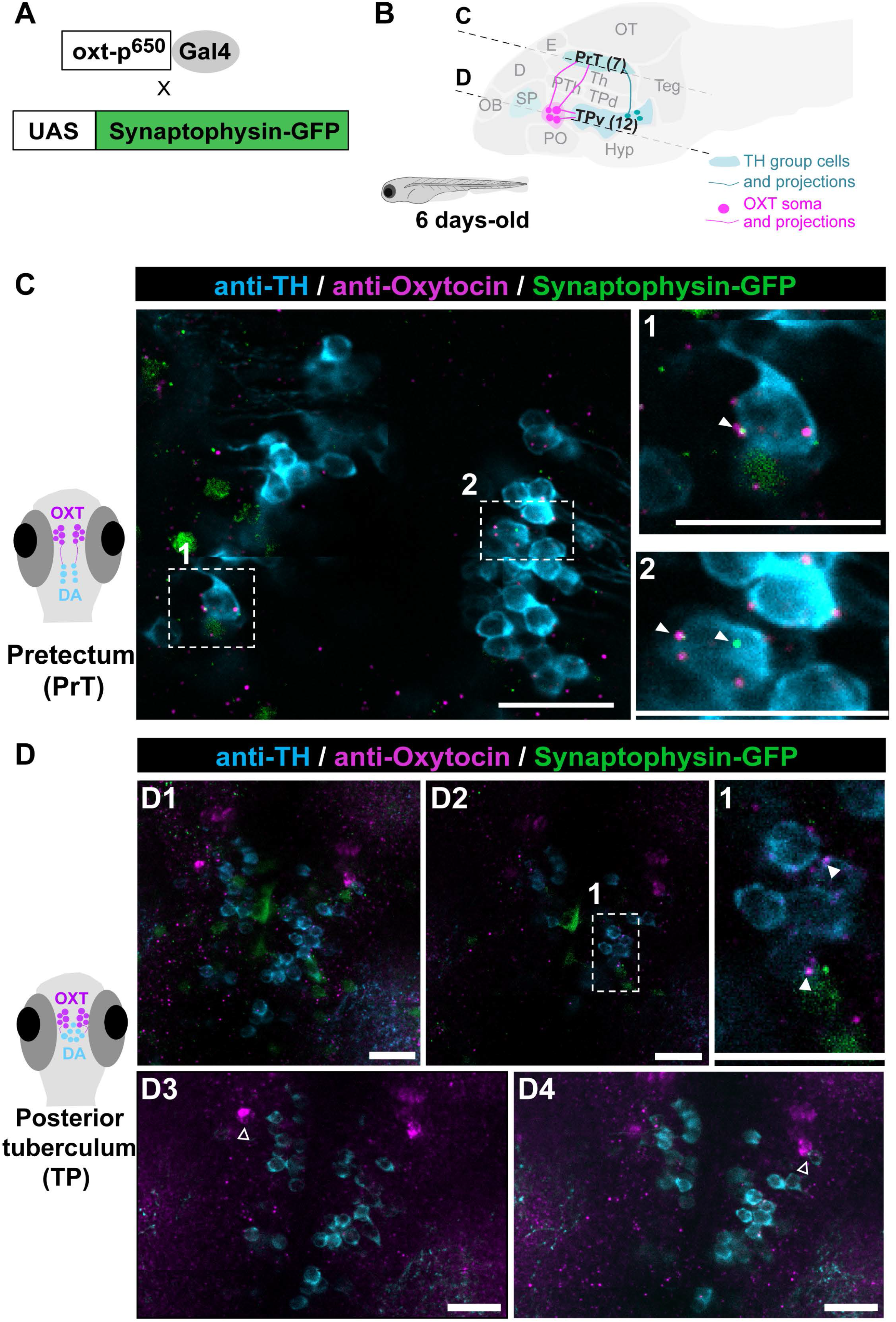
OXT neurons project to pretectal and posterior tuberculum dopaminergic clusters. (A) OXT neuron-specific labeling of synapses: *oxt*:Gal4 drives the expression of the EGFP-fused synaptic vesicle marker synaptophysin-B in oxytocinergic synapses. (B) Schematic of a sagittal larva brain highlighting OXT neuronal projections (in magenta) to pretectal and posterior tuberculum dopaminergic clusters (in blue). (C-D) Representative example showing that the synaptic marker line synaptophysin-GFP (green), driven under the OXT promoter, reveals bona fide OXT synapses containing the OXT peptide (anti-OXT, magenta) on TH positive cells (cyan) in both (C) pretectum (scale: 10 μm) and (D1-D2) posterior tuberculum clusters (scale: 20 μm). (D1) Optical projection overview of the posterior tuberculum. (D2) Single plane image. 1 and 2 depicts amplifications of the OXT synapses on TH positive cells in (C) pretectum and (D2) posterior tuberculum. Closed arrowheads point to OXT positive synaptophysin positive synapses that are in direct contact with TH cells. (D3-D4) 6 slice projection (total 6 μm thickness) of two different planes (scale: 20 μm). Open arrowheads indicate OXT neuron abutting the TH cells of the posterior Tuberculum. Images taken with Zeiss Ism 810 confocal scanning microscope, scaling (x*y*z)= 0.156*0.156*0.96 (µm/pixel), pinhole=31µm, Plan Apochromat 20X/0.8 M27 objective, excitation lasers: green channel (syn-GFP)-488nm; teal (anti-TH) channel-561 nm; magenta (anti-OXT) channel-640 nm).

### Early-life ablation of oxytocinergic neurons leads to impaired social information processing

In view of these results, we next examined whether these long-lasting developmental changes also manifested in the neural processing response to social stimuli in adults, focusing on vertebrate social decision-making network and mesolimbic reward system (O’Connell and Hofmann, 2011). To this end, early-life ablated (MTZ-treated at 4-6 days-old) adult zebrafish were exposed to a single visual social stimulus (a shoal of conspecifics comprised of 2 females and 2 males) or an empty tank for controls, for 10 minutes, and their forebrain neuronal activity state was analyzed by immunostaining of phosphorylated ribosomal protein S6 (pS6), a known correlate for neuronal activation ((Knight et al., 2012), Figure 6). We then quantified the number of pS6-positive neurons in several specific brain areas known to be implicated in neural processing of social information and social reward (O’Connell and Hofmann, 2011) (Figures 6, 6-1, 6-2). Results showed that early OXT ablation significantly affected neuronal activity in response to social stimulus in two specific areas: the anterior part of the parvocellular preoptic nucleus (Ppa) and in the most anterior part of the ventral nucleus of ventral telencephalon (Vv) (GLMM with Poisson regression, planned comparisons and false discovery rate (FDR) corrections, Figures 6B-E, 6-2).

**Figure 6.**
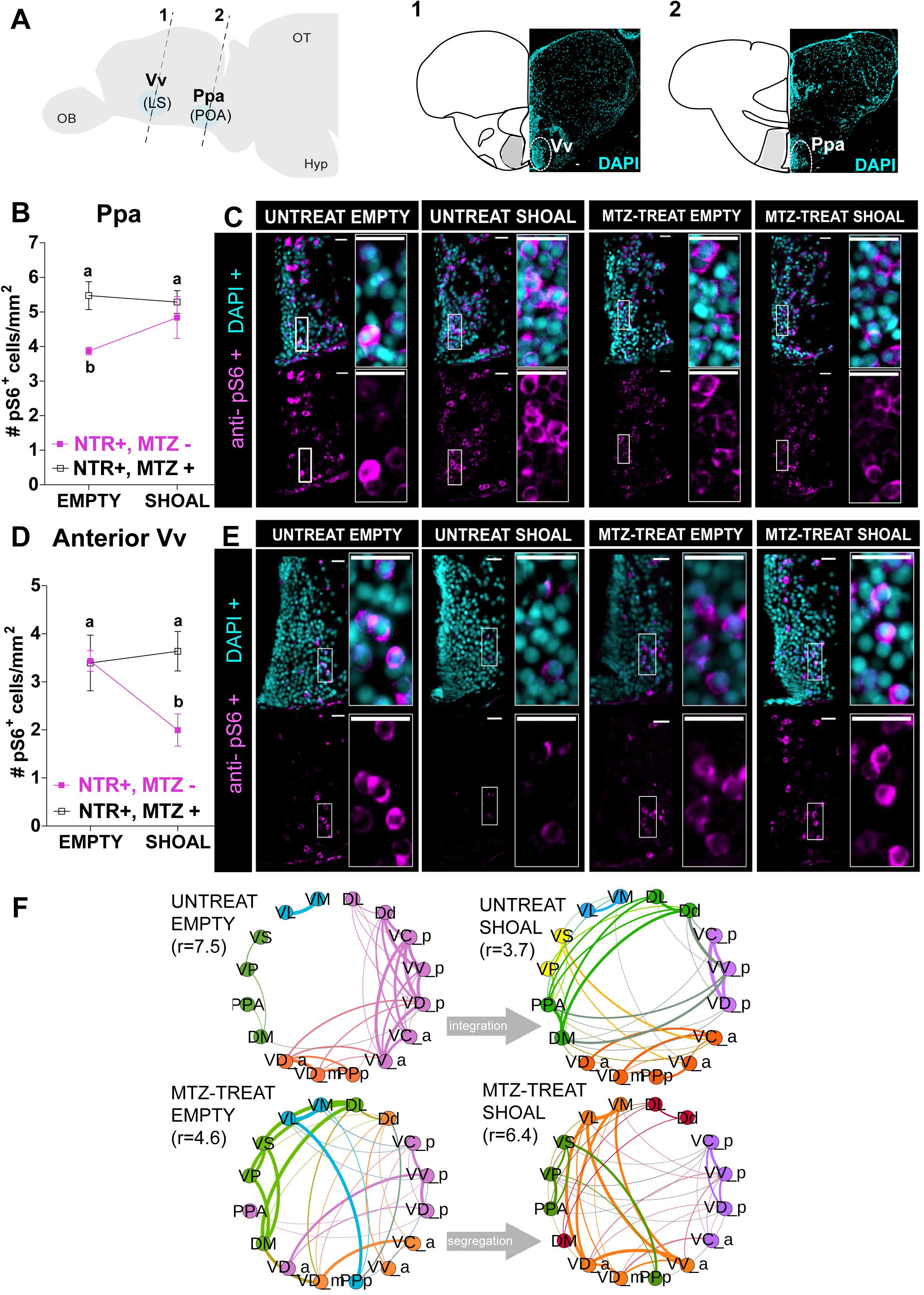
Early-life OXT shapes social information processing. (A) Anatomical localization of the two social responsive areas: anterior part of the ventral nucleus of the ventral telencephalon (Vv_a, 1) and anterior part of the parvocellular preoptic nucleus (Ppa, 2). Brain areas identified by DAPI. (B-E) Quantification of the density (cells/mm^2^) of cells expressing the neuronal marker pS6 in 4-6 days-old MTZ-treated (open squares) or untreated adult fish (full squares), after exposure to either a shoal of conspecifics or an empty tank for 10 min in the Ppa area (B) and anterior Vv area (D) and respective representative examples (C, E). Different letters indicate significant statistical differences *(p* <0.05). (F) Changes in the modular structure of functional connectivity. Modules were obtained by extracting central partition from 400 optimization of Leiden algorithm (Traag et al., 2019) on the treatment correlation matrices. Node color indicates community membership. For visualisation purposes, we only show links with correlation weight >0.1. ***r*** values indicate the ratio of total edge weight within and between modules. High (low) values of ***r*** indicate more (less) segregated modular structure. Scale: 20 μm. Data presented as mean ± SEM.

Specifically, we found that while unablated fish exhibited increased activity in the Ppa upon exposure to a shoal of conspecifics, this was not observed in early ablated fish (untreated shoal vs untreated empty, p= 0.024, n= 11/7 respectively; MTZ-treated shoal vs MTZ-treated empty, p=0.88, n=10/6 respectively; Figure 6B,C, Figure 6-2). Vv neurons of unablated fish exhibited decreased activity upon exposure to a social stimulus, whereas early ablated animals maintained their neuronal activity regardless of the stimulus (untreated shoal vs untreated empty, p= 0.024, n=11/7 respectively; MTZ-treated shoal vs MTZ-treated empty, p=0.88, n= 11/7 respectively; Figure 6D,E; Figure 6-2).

These results show that these two brain areas display deficits in neuronal response to a social stimulus in early OXT-ablated zebrafish. Importantly, the Vv is considered analogous to the mammalian lateral septum (LS), while the Ppa is analogous to the mammalian preoptic hypothalamus (POA) (Wullimann and Mueller, 2004; O’Connell and Hofmann, 2011). These areas are core nodes of the social behavior network in all vertebrate species and have a strong reciprocal connection with each other (Wullimann and Mueller, 2004; O’Connell and Hofmann, 2011).

We next examined whether early-life ablation of OXT neurons in the POA altered functional connectivity between forebrain nuclei belonging to the social decision-making network (SDMN; (O’Connell and Hofmann, 2012)), by comparing the correlation matrices of activity levels for the different areas between ablated and non-ablated fish in either basal state or in response to social stimulus (Figure 6-3, 6-4). We constructed correlation matrices corresponding to each treatment via a resampling scheme, inspired by Quadratic Assignment Procedure (Makagon et al., 2012), that additionally allows to choose the threshold for sparsification on the basis of minimal heterogeneity across resampled correlation matrices (see Methods). To identify functional reconfigurations between different treatments, we identified mesoscale differences in the way the functional networks are organized by their modular (or community) structure. To extract such modules, we performed community detection using the Leiden algorithm (Traag et al., 2019) on the functional connectivity matrix of each treatment. To ensure the robustness of the resulting partitions, we repeated the optimization 400 times per treatment and reconciled the different candidate partitions by considering the central partition (Peixoto, 2021) (see final partitions in Figure 6F). Already by visual inspection it is possible to observe differences in the overall community structure between treatments in the allocation of nodes to communities, and in the relative integration within and across communities. We quantified such integration by calculating the ratio *r* of the total edge weights within communities to the total edge weights between different communities (see Methods). We found that in the basal state (i.e. when no visual stimulus was presented) the mesoscale functional organization is different between ablated (*r* = 4.6) and non-ablated fish (*r* = 7.5), implying that the ablation of OXT neurons in the POA modifies the resting state networks of the forebrain SDMN (Figure 6F). In particular, ablated fish show a much less segregated network with respect to non-ablated ones. Secondly, we observed that, in response to the social stimulus (i.e. sight of a conspecific shoal), the functional network of non-ablated fish transitions from a segregated state (*r* = 7.5) to a more integrated one (*r* = 3.7), whereas ablated fish showed the opposite pattern, moving from a more integrated basal state (*r* = 4.6) to a more segregated response network (*r* = 6.4). All *r* values are significantly different from zero, and also significantly different from each other (*p* < 0.01). Finally, we completed the network analysis focusing on local differences in terms of network centralities. In particular, we looked at the strength of nodes, representing how strongly a node is connected to its neighbors (Figure 6-4). We found a strong and significant anti-correlation between the ranking of strengths for the ablated versus non-ablated fish in response to the social stimulus (Untreated Shoal vs MTZ-treated Shoal, Spearman r=-0.5, p=0.01), implying that the hub nodes for non-ablated fish are poorly connected nodes for the ablated and vice versa. Specifically, in untreated fish exposed to the social stimulus, the most connected nodes were the posterio-ventral part of the ventral telencephalon (Vv_p_), followed by the medial zone of the dorsal telencephalic area (Dm), two regions that have been implicated in the regulation of social interactions and motivated behavior in zebrafish (von Trotha et al., 2014; Stednitz et al., 2018). In contrast, both these nodes drop their centrality in the SDMN network of MTZ-treated fish exposed to the social stimulus (from 1^st^ to 5^th^ and from 2^nd^ to 16^th^, among 16 nodes, respectively), where the anterio-dorsal part of the ventral telencephalon (Vd_a_) and the lateral part of the ventral telencephalon (Vl), two poorly connected nodes in untreated fish (ranking 14^th^ and 16^th^, out of 16 nodes, respectively), were the most connected nodes (Figure 6-4).

These results indicate that early-life OXT ablation leads to blunted response to shoal-induced changes in brain activity within specific nuclei (Vv and Ppa), and to a wide-ranging alteration in network connectivity spanning several nuclei of the social decision-making network in both basal state and in response to social stimulus.

Taken together, our results show that proper response to social stimuli depends on orchestrated co-development of OXT and DA neurons. We show that during development, OXT has important organizational effects. Early developmental perturbation of OXT neurons led to reduced attraction towards conspecifics, impaired the neurodevelopment of specific DA-ergic neurons, caused a blunted neural response to social stimuli in the forebrain (Ppa and Vv), and altered the connectivity of the SDMN.

## DISCUSSION

Previous research has implicated OXT in various developmental disorders that affect social function in humans, have highlighted its importance in social behavior in animals, and indicated the importance of its communication with other systems, such as DA, serotonin and estrogen for proper social behavior (Heinrichs et al., 2009; Hovey et al., 2014; Grinevich et al., 2015; Grinevich and Stoop, 2018; Rajamani et al., 2018). However, there has been limited investigation into the mechanistic aspects of OXTs developmental functions, and how they are linked to its well-described social roles.

Here we utilize temporally controlled perturbations of OXT-ergic cells at different developmental timepoints during zebrafish life to show that OXT has a distinct developmental function, namely to enable proper development of specific downstream DA-ergic clusters known to be involved in visual attention gating and reward (O’Connell and Hofmann, 2011; Antinucci et al., 2019). We also show that in animals where this developmental process was perturbed, the response to social stimuli is affected, and the tendency to shoal with conspecifics is reduced.

The developmentally affected DA-ergic clusters, namely the pretectum (superior colliculus) and the TP (VTA) have been linked in other organisms to social functioning at the level of behavior and, at the molecular level, to the OXT system. Thus, in primates and humans, which like zebrafish are highly dependent on vision for gathering social information, OXT receptors are expressed in the superior colliculus (Schorscher-Petcu et al., 2009; Freeman et al., 2014a, 2014b, 2018), and OXT modulates the gaze and oculomotor responses controlled by this nucleus (Leknes et al., 2013; Kret and De Dreu, 2017). This is also true of the VTA, where OXT receptors are expressed, which functions to promote sociability (Hung et al., 2017). Notably, rhesus monkeys subjected to early social deprivation displayed a reduction in DA neurons of the VTA (Martin et al., 1991), and mouse KO of the autism-associated gene Nlgn3 resulted in impaired OXT signaling in DA-ergic neurons and aberrant activity in the DA-ergic neurons of the VTA (Hörnberg et al., 2020). This further shows how early perturbations to the OXT system can lead to wide-ranging and varied effects on other socially relevant systems in the brain. Whether the changes in DA-ergic neurons, caused the observed changes in social affiliation and neural activity is yet to be directly tested.

In addition to the DA-ergic deficits, we show that in normal fish, neurons of the Ppa (analogous to the mammalian VTA) respond to social stimuli by increased activity, while neurons of the Vv (analogous to the mammalian LS) respond by reducing their activity. In contrast, in developmentally perturbed animals these areas do not change their activity following presentation of the stimulus. Interestingly, the OXTR in the mammalian LS have been implicated in the regulation of social fear (Guzmán et al., 2013; Menon et al., 2018). In zebrafish, the Vv/LS has been functionally associated with social affiliation, mainly through its cholinergic neurons (Stednitz et al., 2018). However, it is interesting to note that in many animal models (e.g. rodents, teleosts, and cartilaginous fish), including zebrafish, the Vv/LS has been shown to contain mainly GABAergic neurons, and to be a source of GABAergic input to the VTA, a putative homologue of the TP, encompassing the TP and PTN, in zebrafish (O’Connell and Hofmann, 2011), and to the POA (O’Connell and Hofmann, 2011; Vega-Quiroga et al., 2018). Thus, it is possible that the observed reduced activity in Vv/LS corresponds to GABAergic neurons that project to the Ppa and TP. These areas are core nodes of the social behavior network in all vertebrate species and have a strong reciprocal connection with each other (Wullimann and Mueller, 2004; O’Connell and Hofmann, 2011). This suggests that social stimuli could promote a disinhibition in these target areas, which would be associated with the rewarding and/or motivational component of social affiliation. As a corollary, in early-ablated animals, where Vv activity remains high in response to social stimuli, the TP would remain inhibited, and the rewarding component of social affiliation would be attenuated, leading to decreased display of this behavior.

In addition, we show that the overall coactivation patterns of forebrain nuclei involved in social processing are altered in developmentally perturbed animals, both at baseline and following presentation of a social stimulus. In other words, we show that the connectivity between the different nodes of the social decision-making network changes as a result of early life OXT ablation. We submit that the proper maturation of this network at the level of neural architecture is dependent on early-life signaling by OXT neurons.

In summary, our results demonstrate that OXT neurons are required for the developmental acquisition of social affiliation and exert an organizational effect on DA-ergic neuronal populations. Furthermore, our results suggest that this role involves orchestrating the co-development and maturation of several brain systems, such as the midbrain social visual processing and attention systems (the pretectum), social decision-making areas in the forebrain and ascending dopamine centers associated with reward. When the OXT-dependent developmental process is perturbed, neural responses to social stimuli in these regions become compromised, important parts of the social decision-making network fail to synchronize their activity, and eventually, the propensity to spend time near a shoal of conspecifics is reduced.

## ACKNOWLEDGMENTS

We thank all members of Gil Levkowitz’s and Rui Oliveira’s laboratories for fruitful discussions, and Nitzan Konstantin for English editing. We thank the technical support of IGC’s Advanced Imaging Facility (AIF-UIC), which is supported by the national Portuguese funding ref# PPBI-POCI-01-0145-FEDER-022122, co-financed by Lisboa Regional Operational Programme (Lisboa 2020), under the Portugal 2020 Partnership Agreement, through the European Regional Development Fund (FEDER) and Fundação para a Ciência e Tecnologia (FCT, Portugal); all the staff from the Fish Facility platforms of Instituto Gulbenkian de Ciência, Portugal, and Weizmann Institute of Science, Israel, for animal care and valuable advice; Congento, which is supported by the funding ref# LISBOA-01-0145-FEDER-022170, co-financed by Lisboa Regional Operational Programme (Lisboa 2020), under the Portugal 2020 Partnership Agreement, through the European Regional Development Fund (FEDER) and Fundação para a Ciência e a Tecnologia (FCT; Portugal); IGC’s Histopathology Facility for technical support, help and valuable advices; the Advanced BioImaging and BioOptics Experimental Platform (ABBE Platform) of the Champalimaud Center for the Unknown (CCU) for all the technical support and help.

## Authors contributions

ARN-Designed the research, performed experiments, analyzed data, wrote/revised manuscript; MG - Designed the research, performed experiments, analyzed data, wrote/revised manuscript; EW- Performed experiments; JB- Performed experiments; GP and MT- Performed network analysis; SAMV- Performed statistical analysis; GL- Designed the research, supervision, edited/revised manuscript; RO- Designed the research, supervision, edited/revised manuscript

---

**Figure 2-1:**
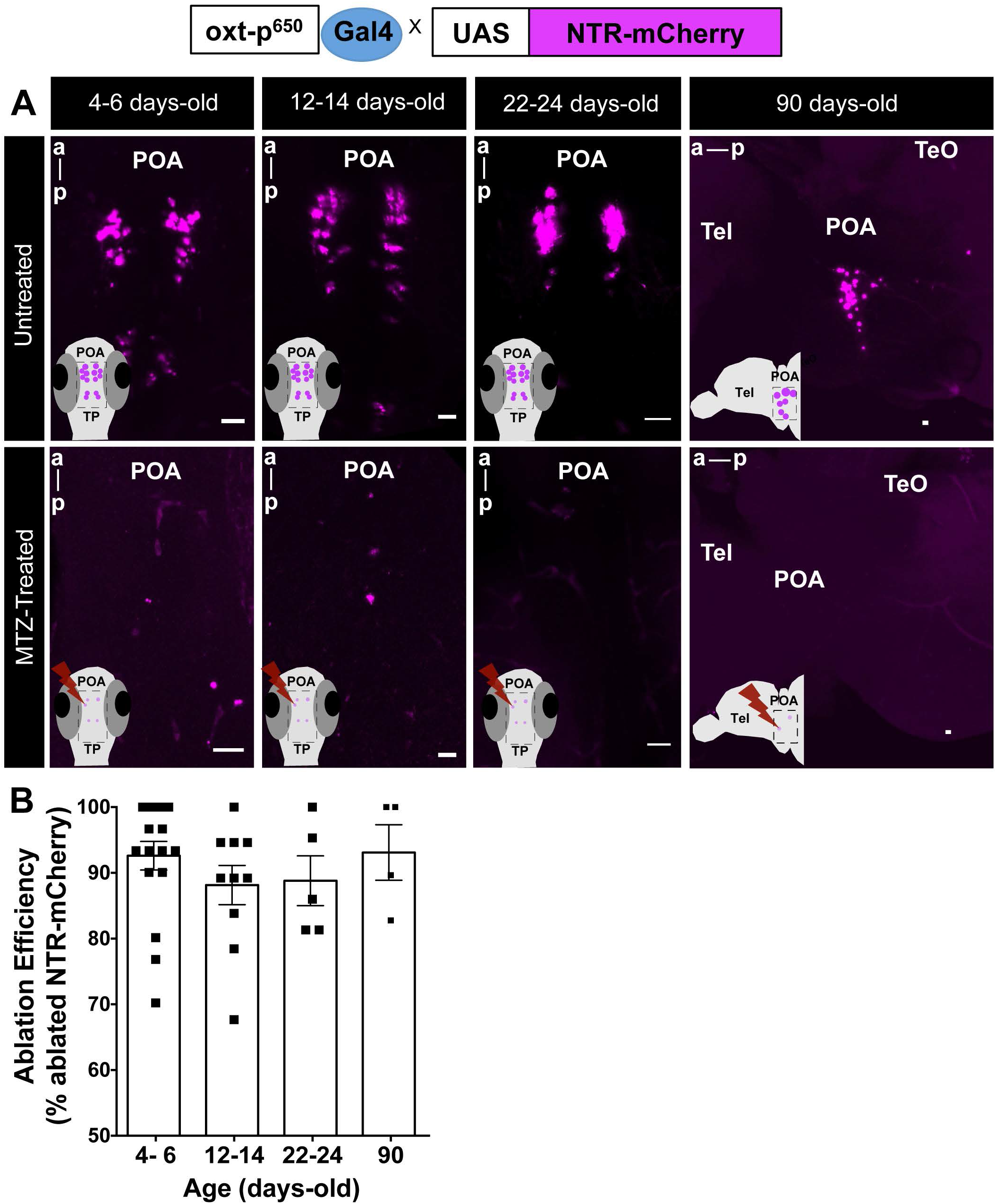
Spatio-temporal control of OXT-specific transgene expression. (A) Representative example of the MTZ-treatment effect on transgene (*oxt:GAL4/UAS-NTR-mCherry*) at different treatment time-points. For 4-6, 12-14 and 22-24 days-old: whole mount larvae, maximum intensity confocal z-stack image, dorsal view, anterior to top; for 90 days-old: sagittal brain slice, maximum intensity confocal z-stack image, anterior to left. Scale: 20 µ*m*. (B) Quantification of the effect of MTZ treatment on transgene expression at all ages tested (4-6 days-old, n=17; 12-14, n=10; 22-24 days-old, n=5; 90 days-old, n=4). Ablation efficiency was measured as percentage of ablated cells in MTZ-treated fish over mean number of NTR-mcherry cells in untreated fish. *POA*, neurosecretory preoptic area; *Tel*, Telencephalon; *TeO*, tectum optic.

**Figure 2-2:**
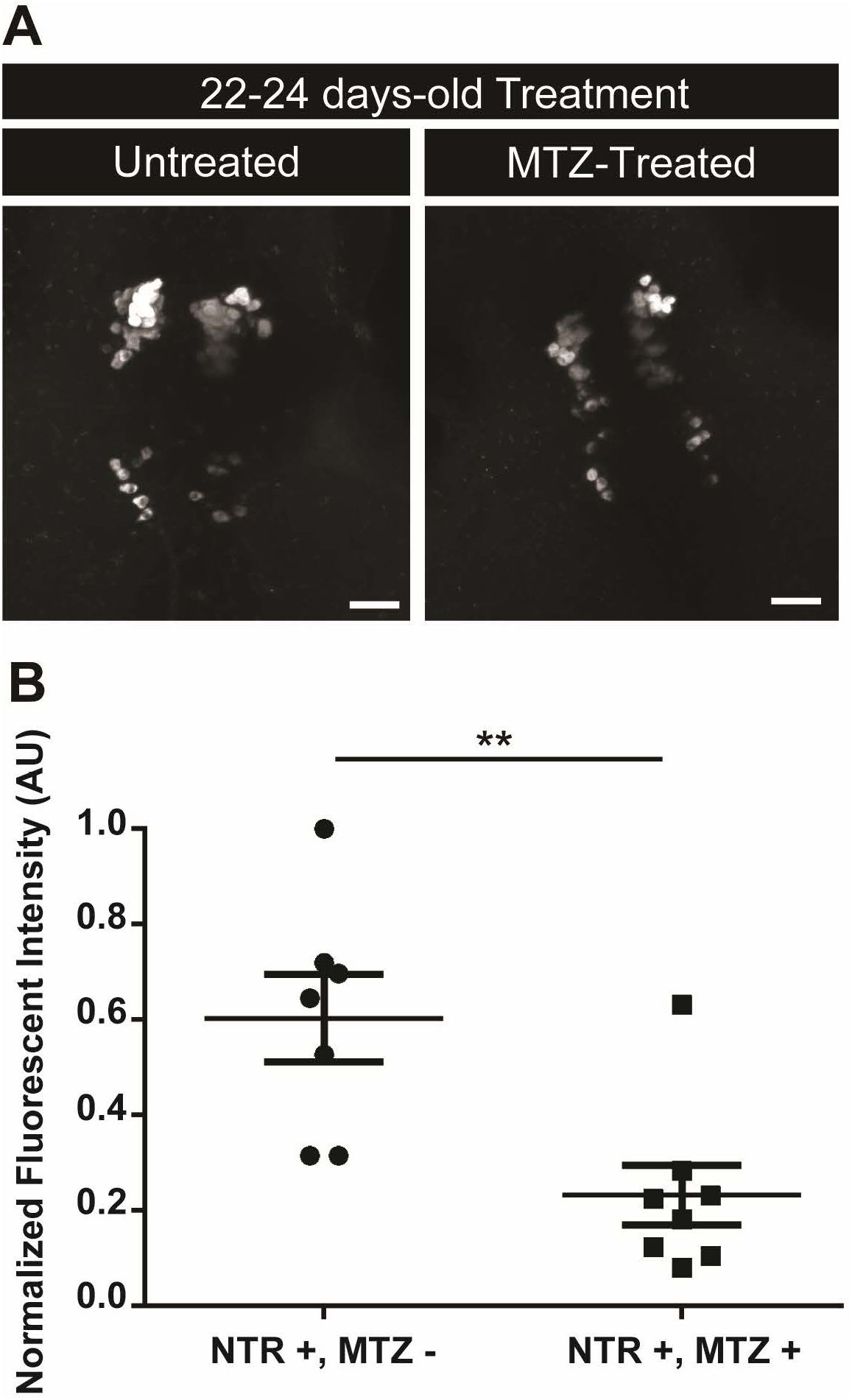
Effects of metronidazole (MTZ)-induced ablation on endogenous OXT at 22-24 days-old treatment by in situ hybridization. (A) Representative example of the 22-24 days-old MTZ-treatment effect on endogenous OXT. Whole mount larvae, maximum intensity confocal z-stack image, dorsal view, anterior to top. Scale: 20 µ*m*. (B) Quantification of normalized fluorescent intensity (AU) in 22-24 days-old untreated (NTR+, MTZ-) or MTZ-treated fish (NTR+, MTZ+). Data presented as mean ± SEM. **p<0.01

**Figure 2-3.**
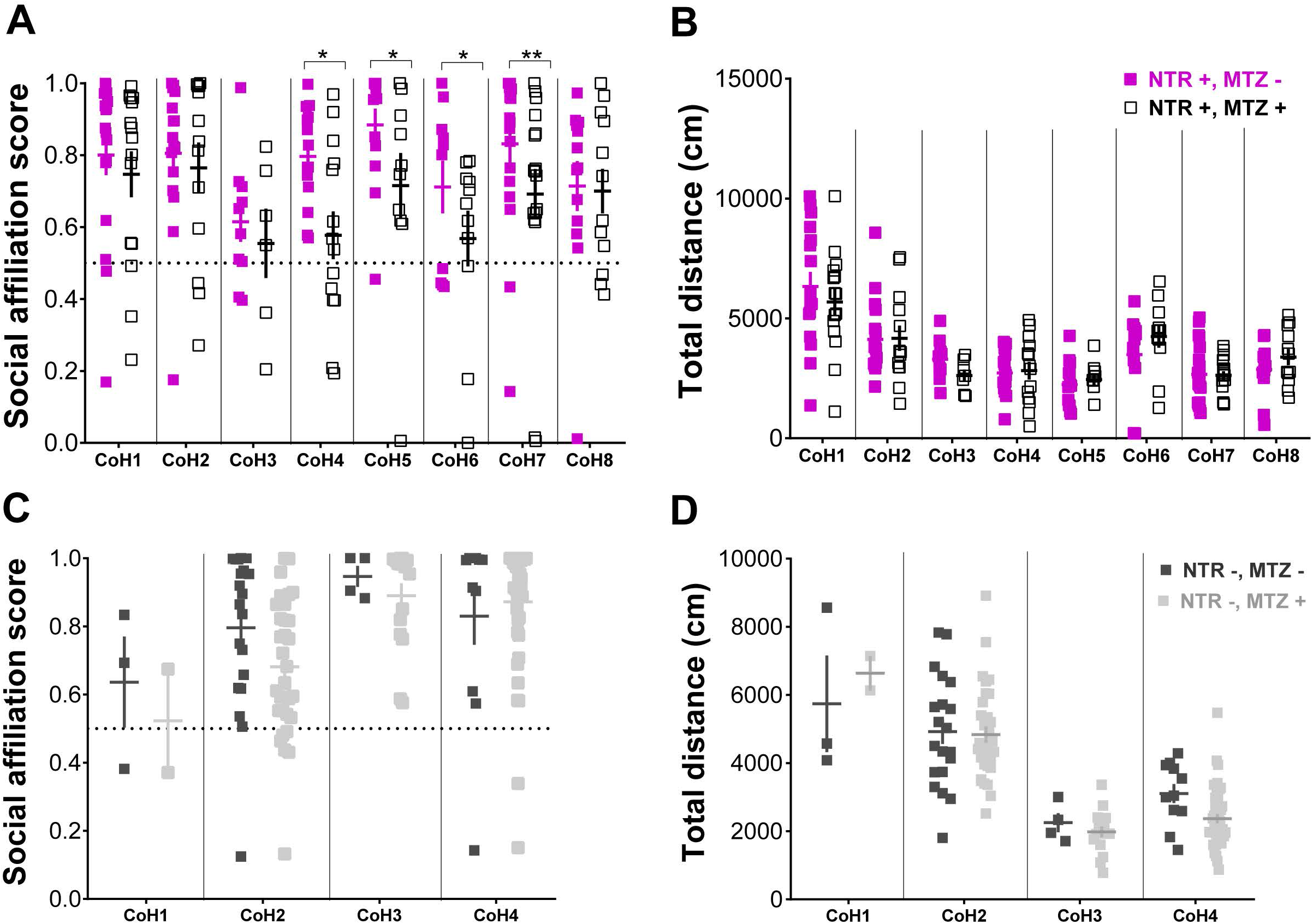
Organizational vs activational effects of oxytocin neurons in adult social affiliation. Effect of early (4-6 days-old) MTZ treatment on (A) adult social affiliation and total distance moved of eight independent cohorts of either MTZ treated at 4-6 days-old (NTR+, MTZ+) or untreated control fish (NTR+, MTZ-); (C) adult social affiliation and (D) total distance moved of control cohorts not expressing the transgene, 4-6 days-old MTZ-treated (NTR-, MTZ+) or untreated (NTR-, MTZ-). One-tailed p-values were considered in (A) because of our *a priori* directionality hypothesis that by ablating OXT neurons, MTZ treatment of fish expressing NTR transgene would decrease social affiliation behavior. Data are presented as mean ± SEM. Full squares (purple or dark gray): untreated fish (MTZ-); open square or light gray: MTZ-treated fish (MTZ+).

**Figure 2-4.**
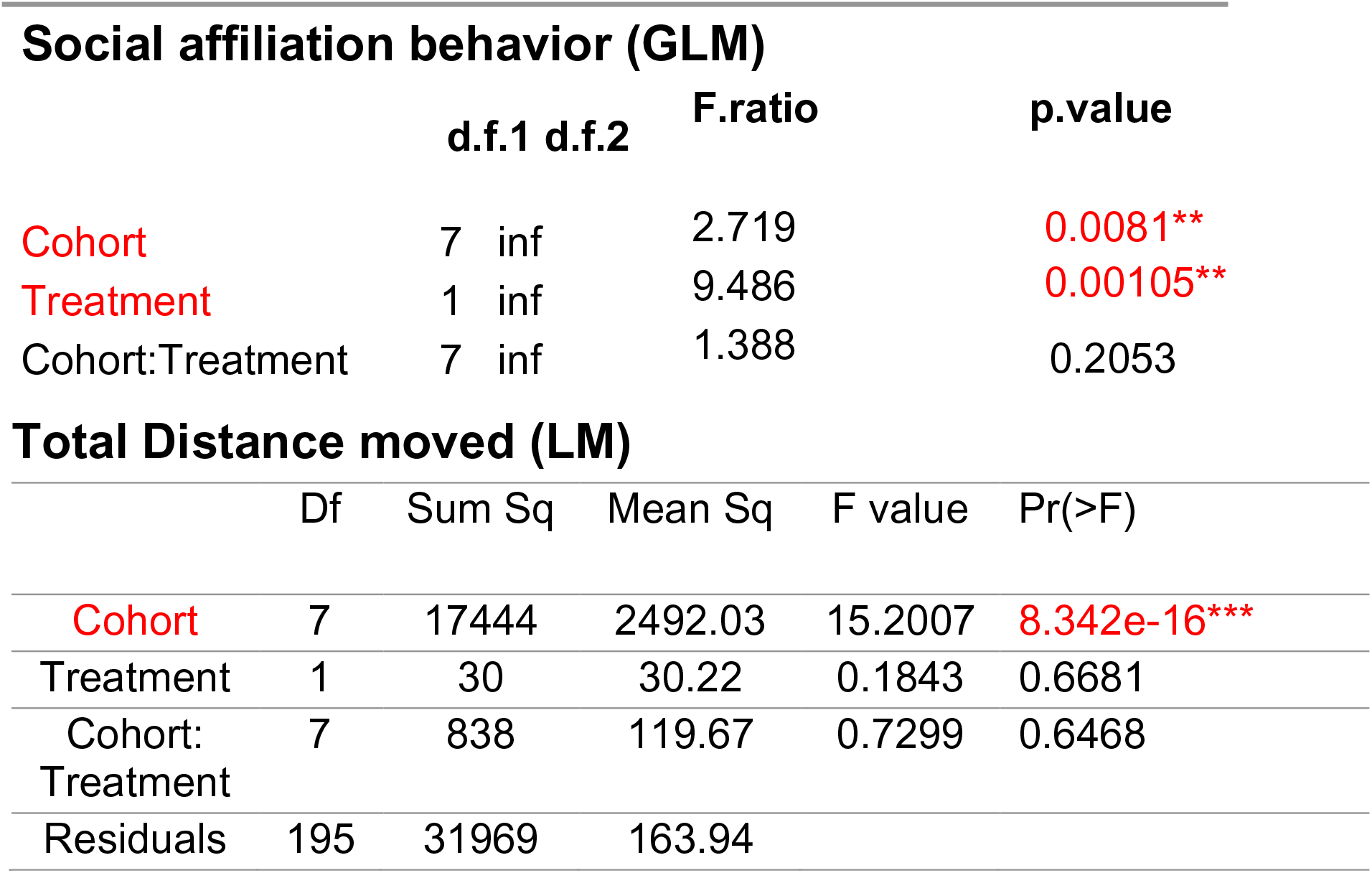
Effects of 4-6 days-old MTZ treatment in adult social affiliation and total distance moved on eight independent experimental cohorts tested. Summary of results of the GLM (Generalized Linear Model) with beta regression and LM (Linear Model) models to analyse the effects of early (4-6 days-old) MTZ-treatment on adult social affiliation and total distance moved on eight independent experimental cohorts expressing the NTR transgene. A one-tailed p-value was considered for the effect of MTZ-treatment on social affiliation behaviour because of our a priori directionality hypothesis (see Methods/Statistic section).

**Figure 2-5.**
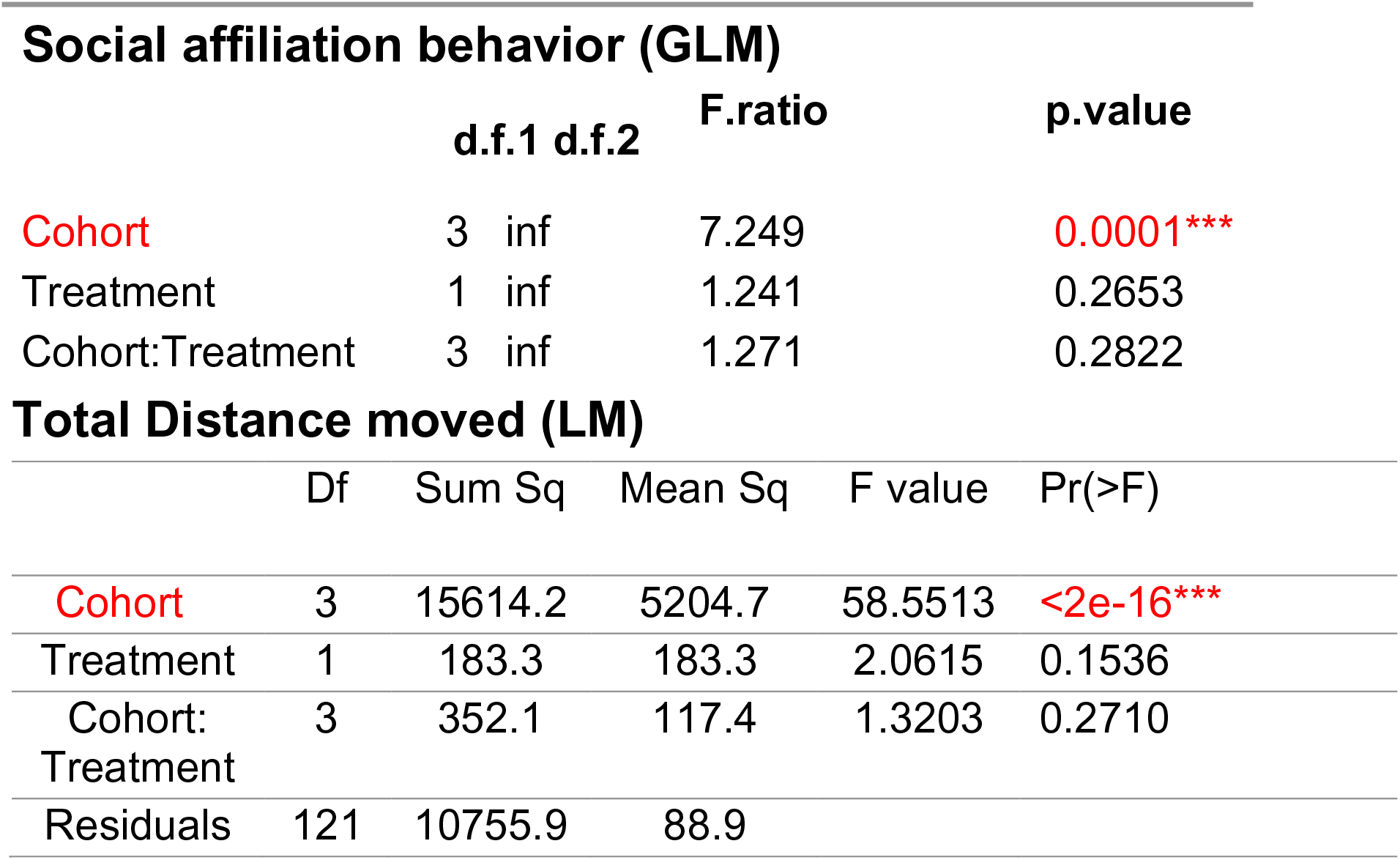
Effects of 4-6 days-old MTZ-treatment in adult social affiliation and total distance moved on control fish not expressing the NTR transgene. Summary of results of the GLM with beta regression and LM models to analyse the effects of early (4-6 days-old) MTZ-treatment on adult social affiliation and total distance moved on control fish not expressing the NTR transgene.

**Figure 2-6.**
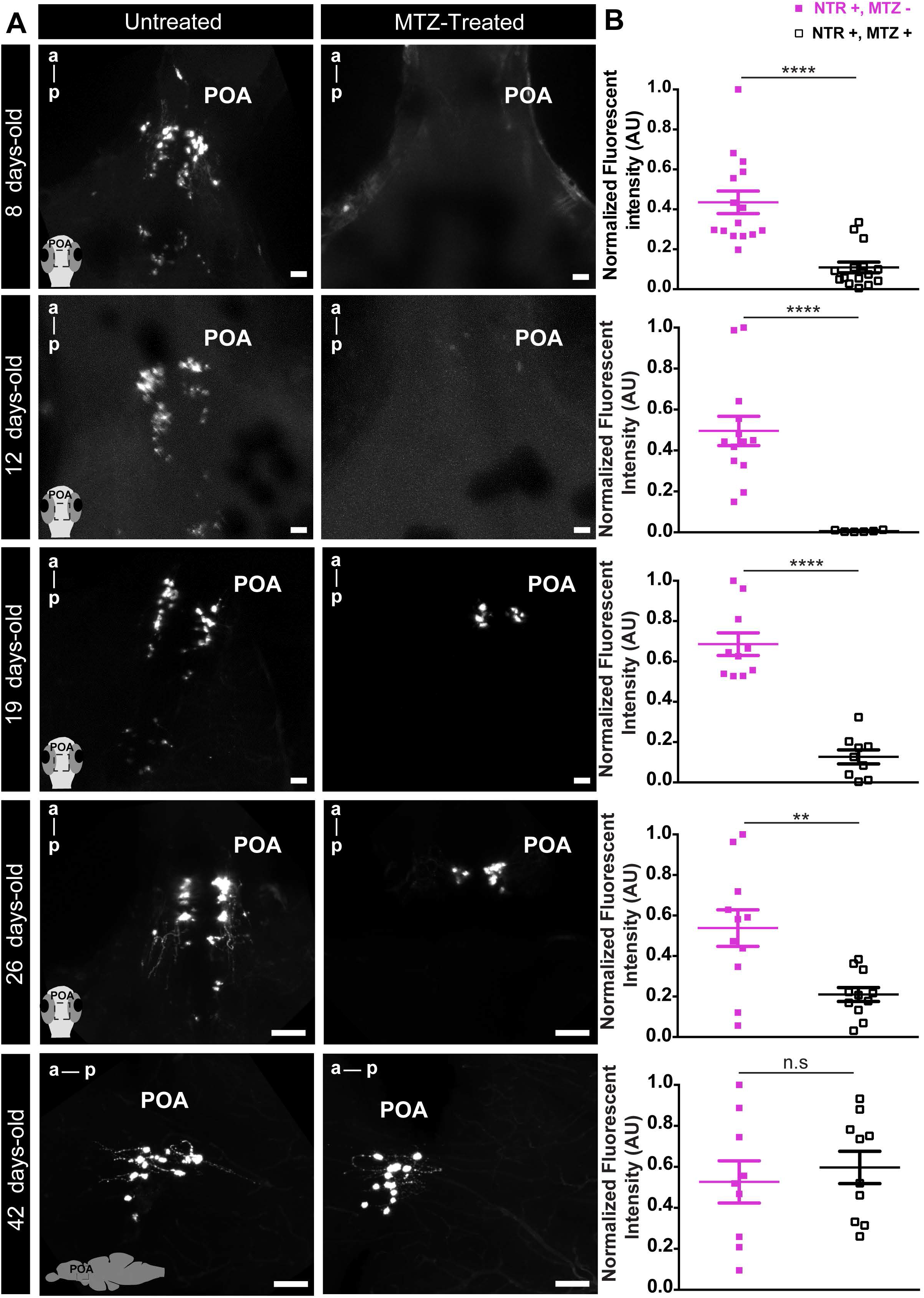
Time-course OXT:NTR-mCherry cell recovery after 4-6 days-old MTZ-treatment. 4-6 days-old metronidazole (MTZ)-treated *vs* untreated larvae were left to grow and sampled at different time-points throughout development: 8, 12, 19, 26, and 42 days-old. A) Representative whole mount larvae, maximum intensity confocal z-stack image, dorsal view, anterior to top (8, 12, 19 and 26 days-old) and sagittal brain slice, confocal z-stack image, anterior to left (42 days-old). B) Quantification of the Normalized fluorescent intensity (AU) in untreated (full squares) versus MTZ-treated fish (open squares) at the different time points sampled. Scale: 20 µ*m* (8-12-19-days-old) and 50 µ*m* (26 and 42 days-old). Data are presented as mean ± SEM. Full squares: untreated fish (NTR+,MTZ-); open square: MTZ-treated fish (NTR+,MTZ+). **p<0.01,**** p<0.0001

**Figure 2-7.**
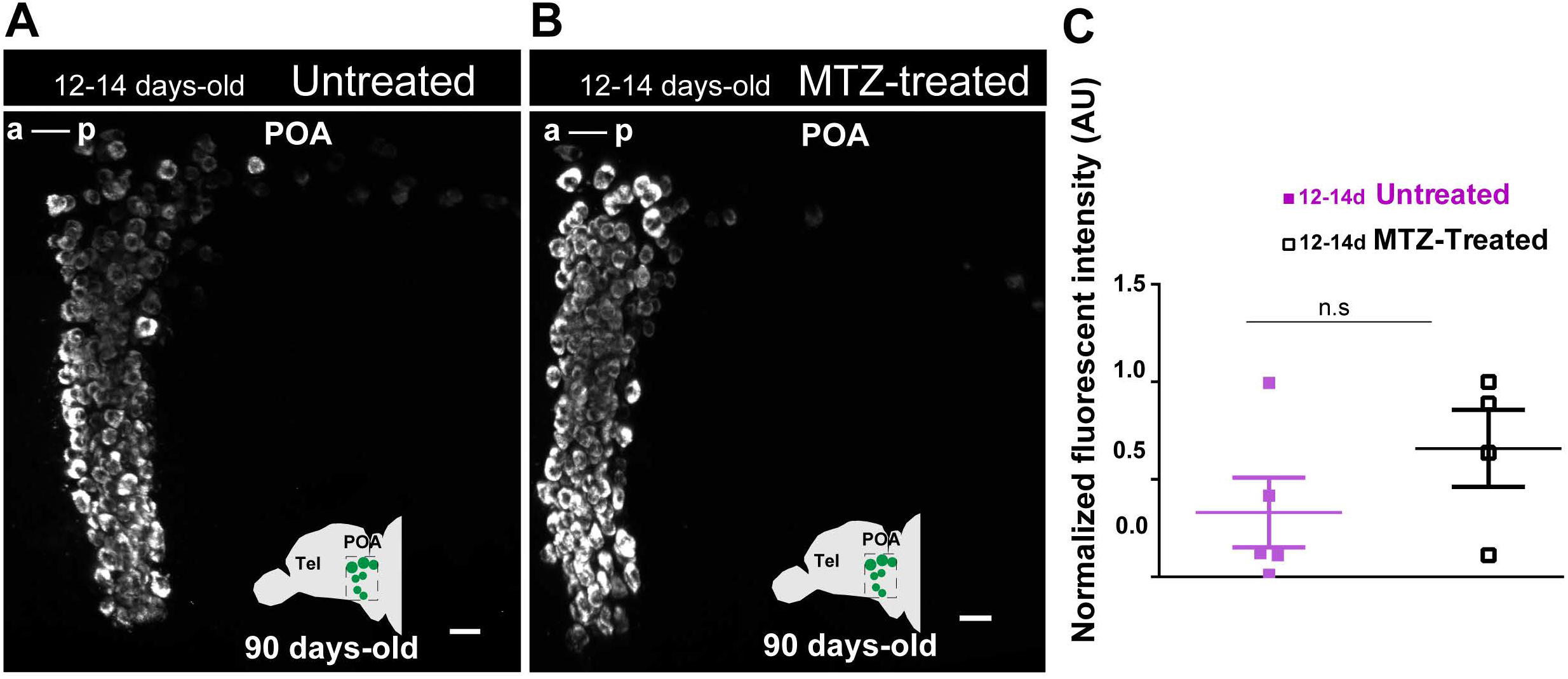
Endogenous OXT mRNA recovery in adulthood after 12-14 days-old MTZ-treatment. (A) Representative example of the endogenous OXT mRNA in untreated adult fish. (B) Representative example of the endogenous OXT mRNA in adult fish treated with MTZ at 12-14 days-old. Images in (A,B) are maximum intensity confocal z-stacks, sagittal slices, anterior to left. Scale: 20 µ*m*. (C) Quantification of normalized fluorescent intensity (AU) in untreated (full squares) and 12-14 days-old MTZ-treated adult fish (empty squares). Data presented as mean ± SEM. Full squares: untreated fish (NTR+,MTZ-); open square: MTZ-treated fish (NTR+,MTZ+).

**Figure 3-1.**
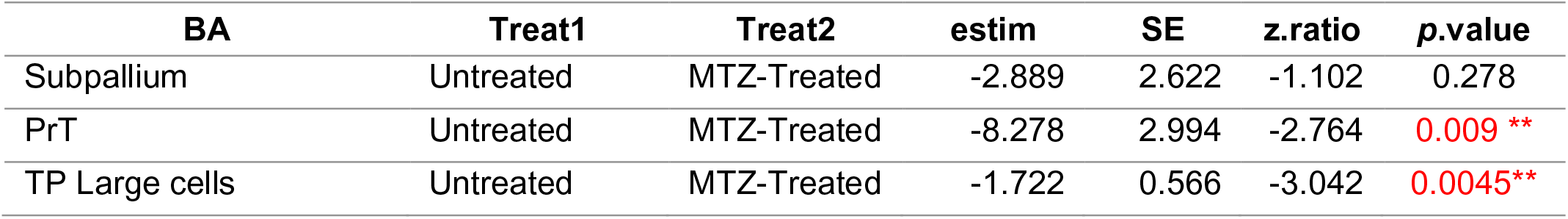
Effect of 4-6 days-old MTZ-treatment in TH cell clusters of larvae zebrafish brain. Three independent brain areas were analyzed. Linear model with post-hoc tests comparing the treatment (early MTZ-treated vs. untreated) were performed. *BA*, Brain area; *Treat*, Treatment; *estim*, estimate; *PrT,* Pretectum; *TP*, Posterior Tuberculum.

**Figure 4-1.**
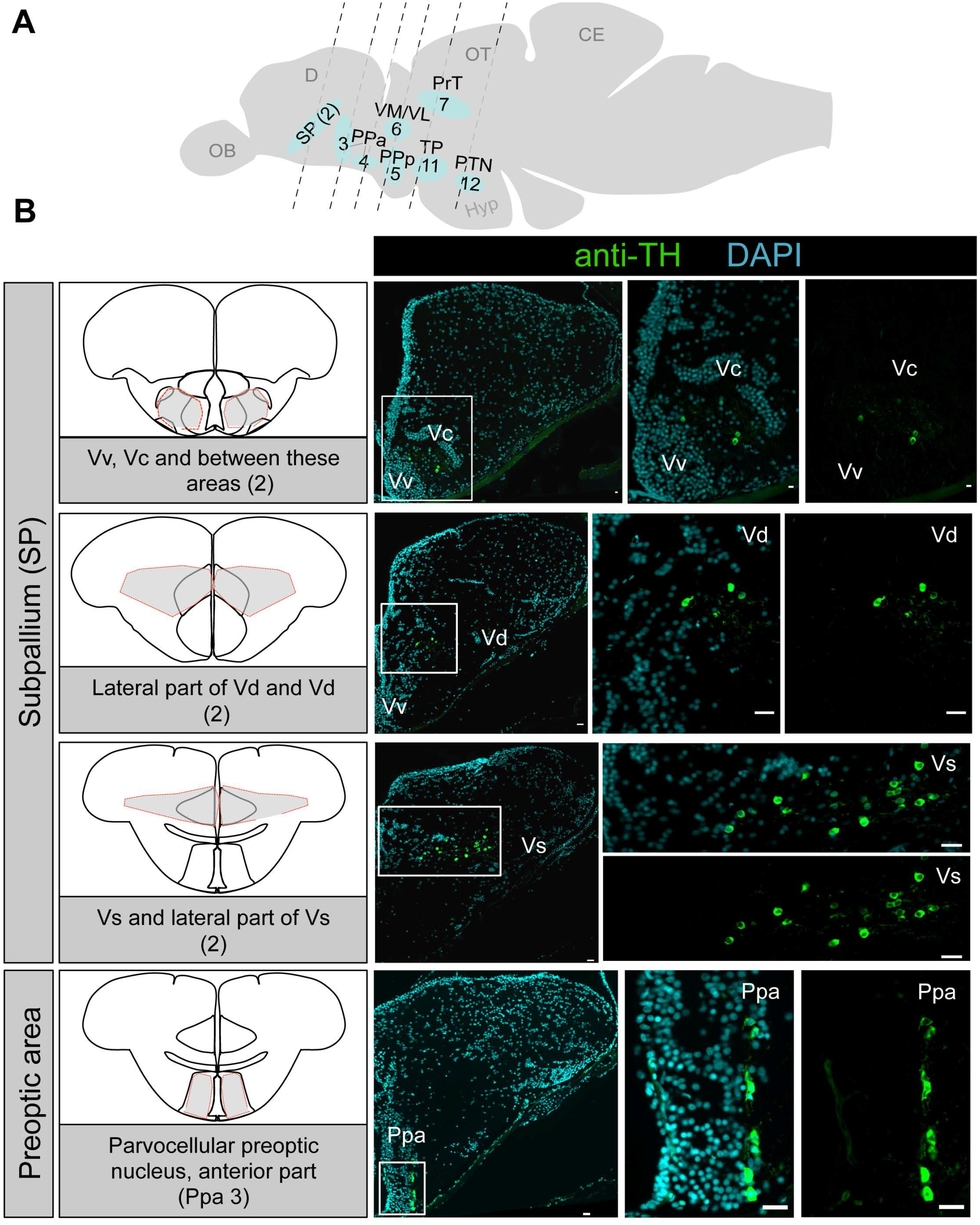
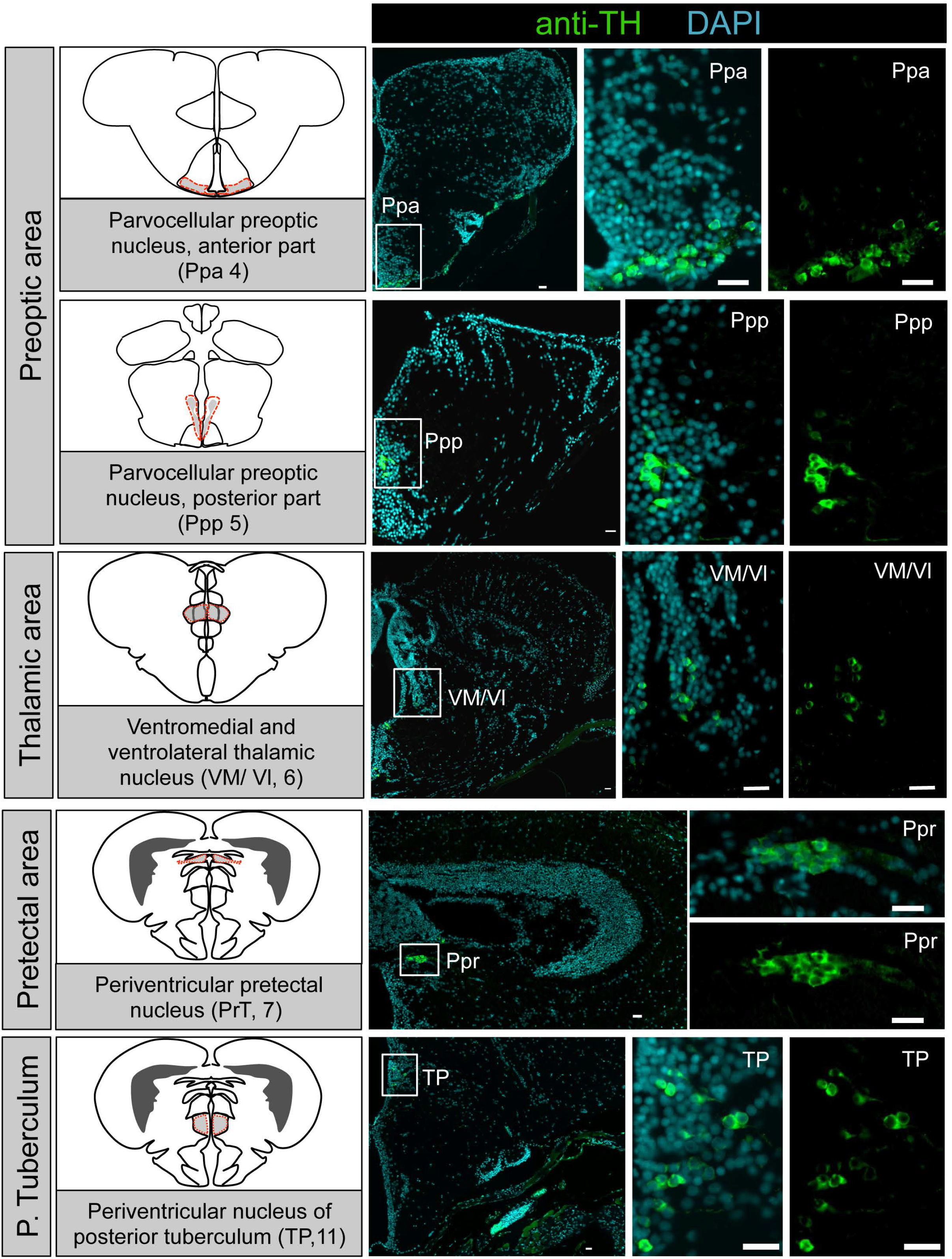
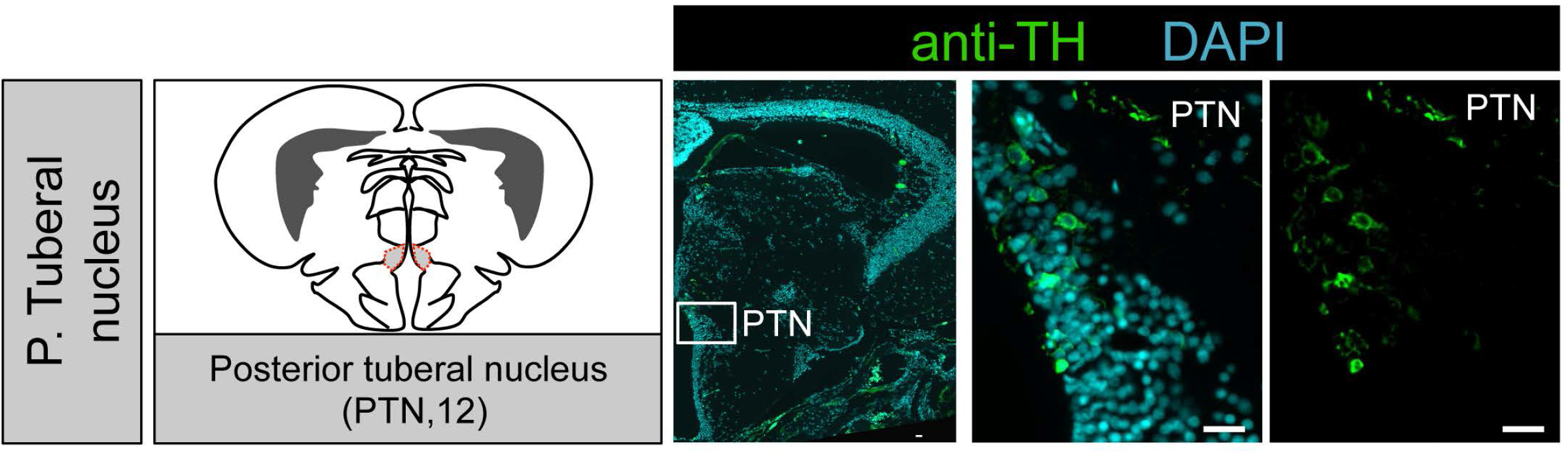
Early life OXT ablation affects the dopaminergic system. (A) Schematic representation of an adult zebrafish sagittal view representing the coronal sections of all dopaminergic areas sampled in the adult brain (dopaminergic cluster nomenclature according to(Panula et al., 2010)). (B) Schematic representation of zebrafish brain coronal sections highlighting the different dopaminergic clusters (adapted from (Wullimann et al., 1996)) and representative example showing anatomical localization of the dopaminergic groups in an untreated adult zebrafish. Landmarks of the areas identified by DAPI (cyan) and dopaminergic groups by TH (Tyrosine Hydroxylase) immunostaining (green). Scale bars are 20μm.

**Figure 4-2.**
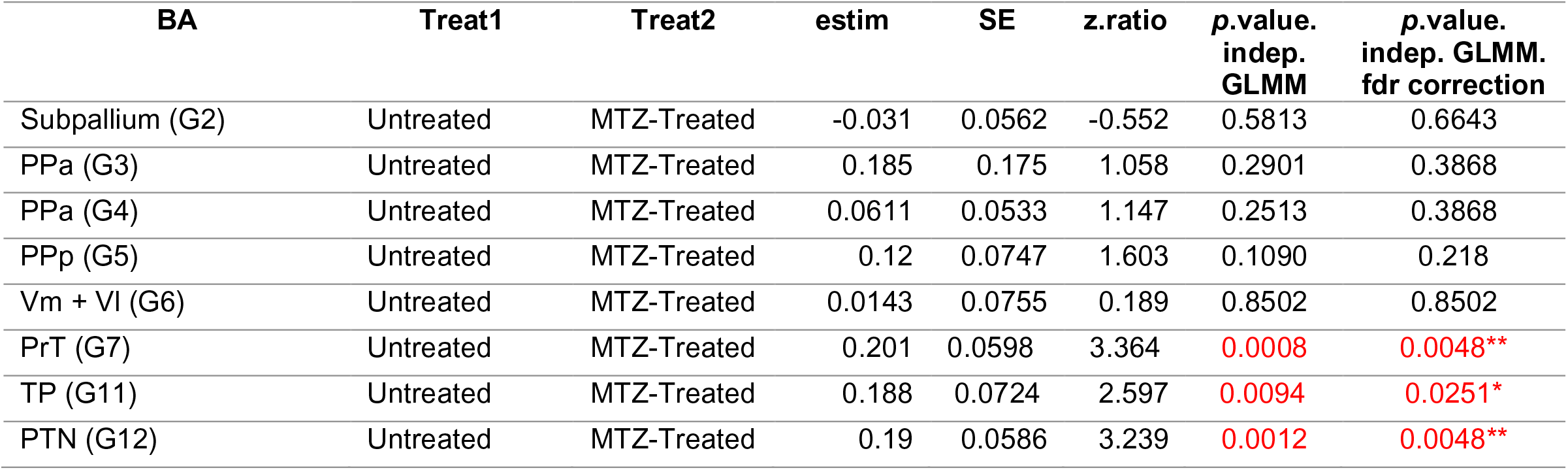
Effect of 4-6 days-old MTZ-treatment in TH cell clusters of adult zebrafish brain. Eight independent brain areas were analyzed. GLMM with a Poisson regression and planned comparisons (early MTZ-Treated vs. Untreated) was performed. *p*-values from planned comparisons were corrected with false discovery rate (FDR) adjustment method. *BA*, Brain area; *Treat*, Treatment; *estim*, estimate; *GLMM*, Generalized Linear Mixed Models; *FDR*, false discovery rate, *PPa*, anterior part of the parvocellular preoptic nucleus; *PPp*, posterior part of the Parvocellular preoptic nucleus; *Vm*, ventromedial thalamic nucleus; *Vl*, ventrolateral thalamic nucleus; *PrT*, Pretectum nucleus; *TP*, Posterior tuberculum nucleus; *PTN*, Posterior Tuberal nucleus.

**Figure 6-1.**
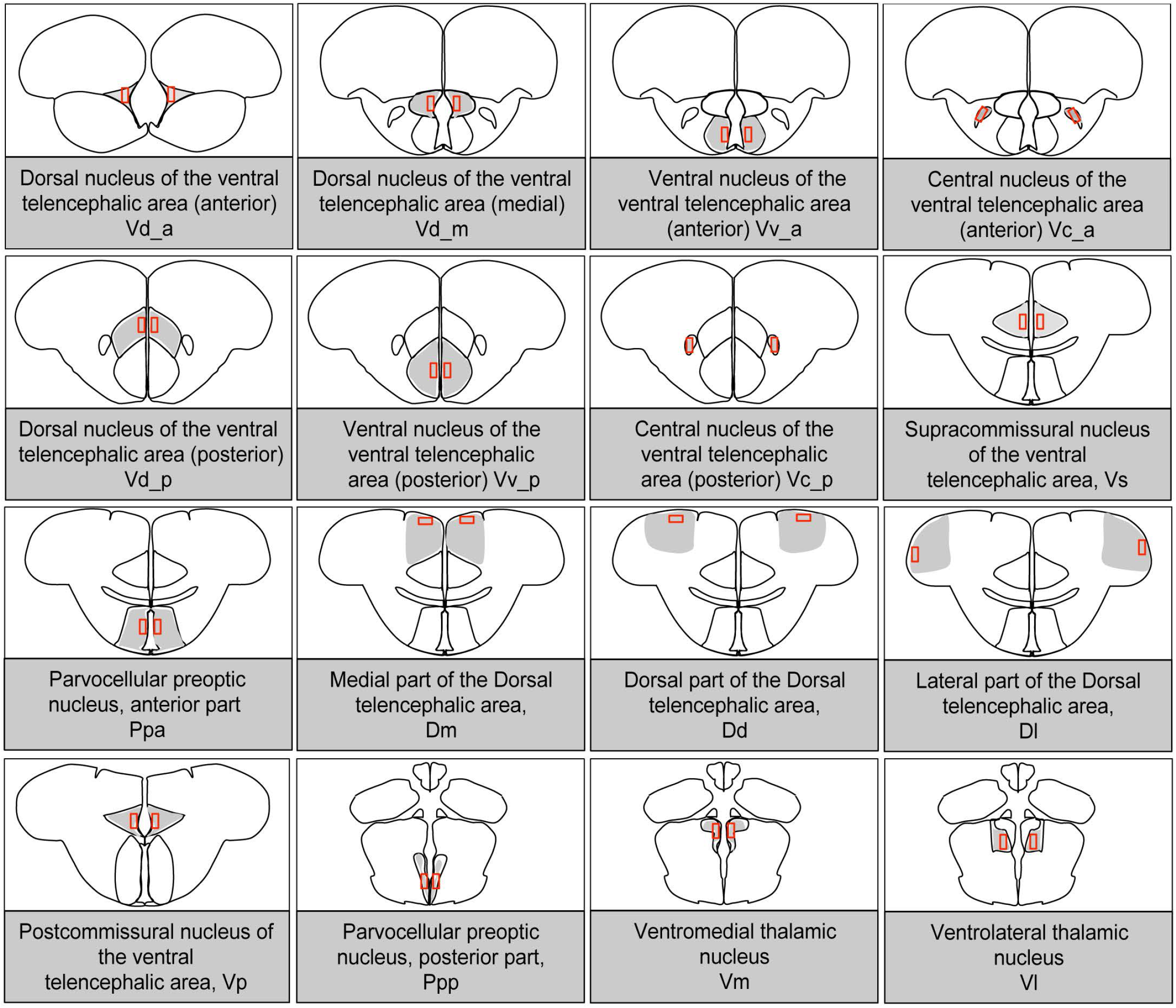
Schematic representation of brain areas that were analyzed for neuronal activation in response to social stimulus. pS6 immunostaining served as a readout of neuronal activation following MTZ treatment at 4-6 days-old or untreated adult fish to either a shoal of conspecifics or an empty tank. Included brain areas are part of the social decision-making network, subdivided into more anterior or posterior parts. Schematics are adapted from zebrafish atlas (Wullimann et al., 1996).

**Figure 6-2.**
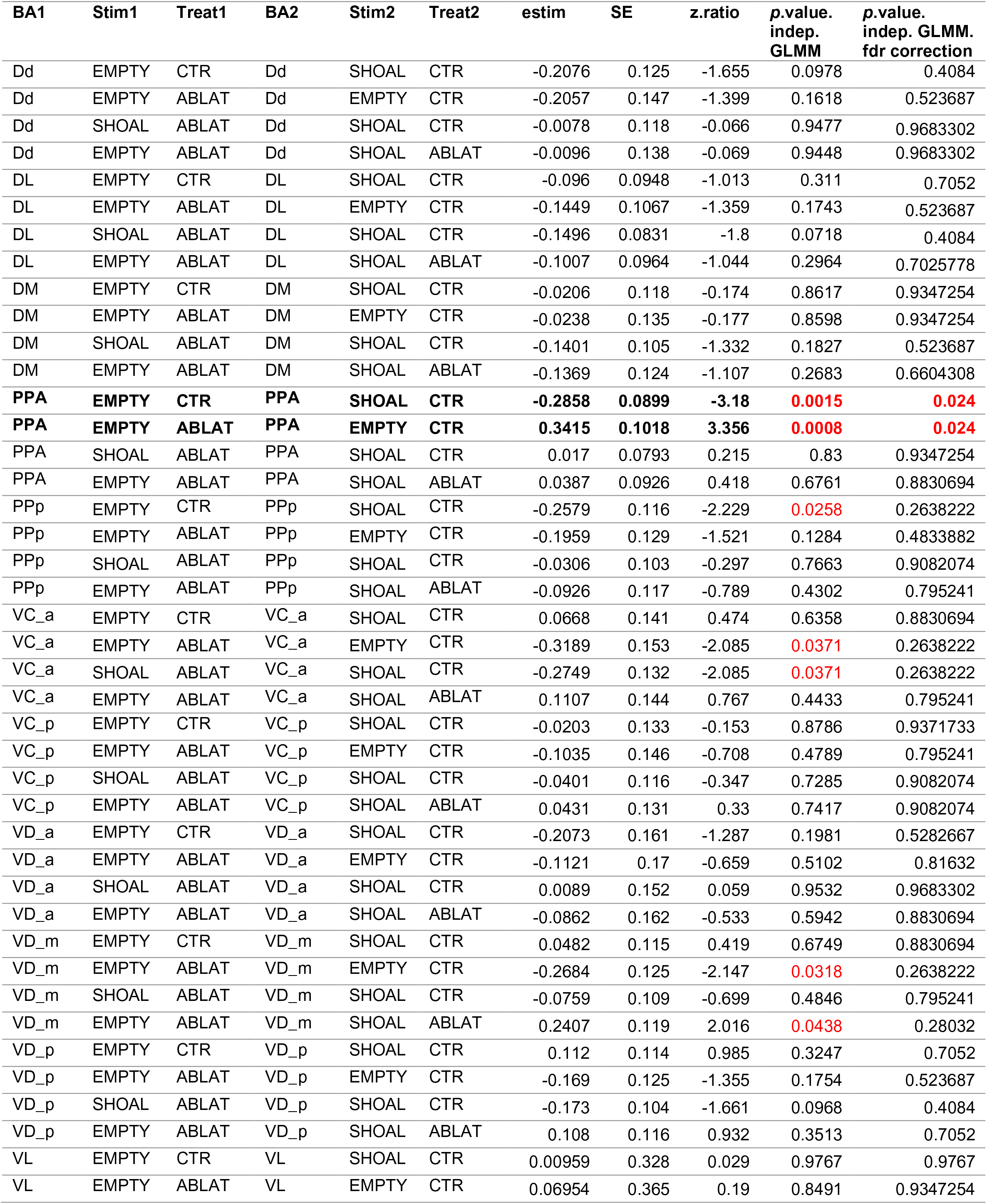
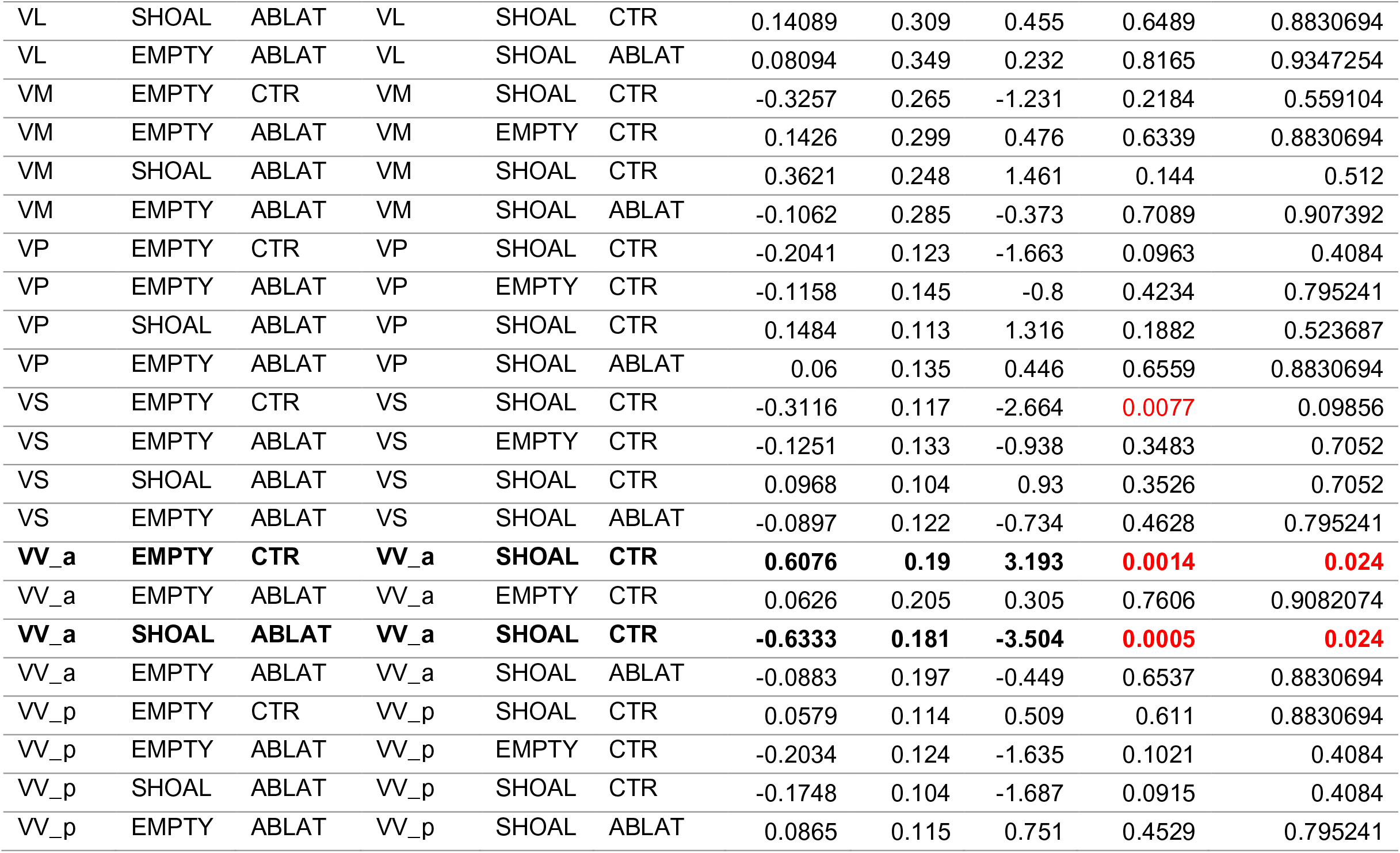
Effect of 4-6 days-old MTZ-treatment on adult neuronal activation. Neuronal activation measured by pS6 immunostaining. Sixteen (16) independent brain areas were analyzed. GLMM with a Poisson regression and planned comparisons were performed. *p*-values of the planned comparisons were corrected with false discovery rate (FDR) adjustment method (significant values in red). *BA1*, Brain area 1; *BA2*, Brain area 2; *Stim*, stimulus; *Treat*, Treatment; *estim*, estimate; *indep*., independent, *GLMM*, Generalized Linear Mixed Models; *FDR*, false discovery rate, *CTR*, untreated fish; *ABLT*, early MTZ-treated fish; *Vv_a*, anterior part of the ventral nucleus of the ventral telencephalic area (V); *Vv_p*, posterior part of the ventral nucleus of the ventral telencephalic area (V); *Vd_a*, anterior part of the dorsal nucleus of V; *Vd_m*, medial part of the dorsal nucleus of V; *Vd_p*, posterior part of the dorsal nucleus of V; *Vc_a*, anterior part of the central nucleus of V; *Vc_p*, posterior part of the central nucleus of V; *Dm*, medial zone of the dorsal telencephalic area (D); *Dl*, lateral zone of the dorsal telencephalic area (D); *Dd*, dorsal zone of the dorsal telencephalic area (D); *Vs*, supracommissural nucleus of V; *Vp*, postcommissural nucleus of V; *Ppa*, anterior part of the parvocellular preoptic nucleus; *Ppp*, posterior part of the parvocellular preoptic nucleus; *VM*, ventromedial thalamic nucleus; *VL*, ventrolateral thalamic nucleus.

**Figure 6-3.**
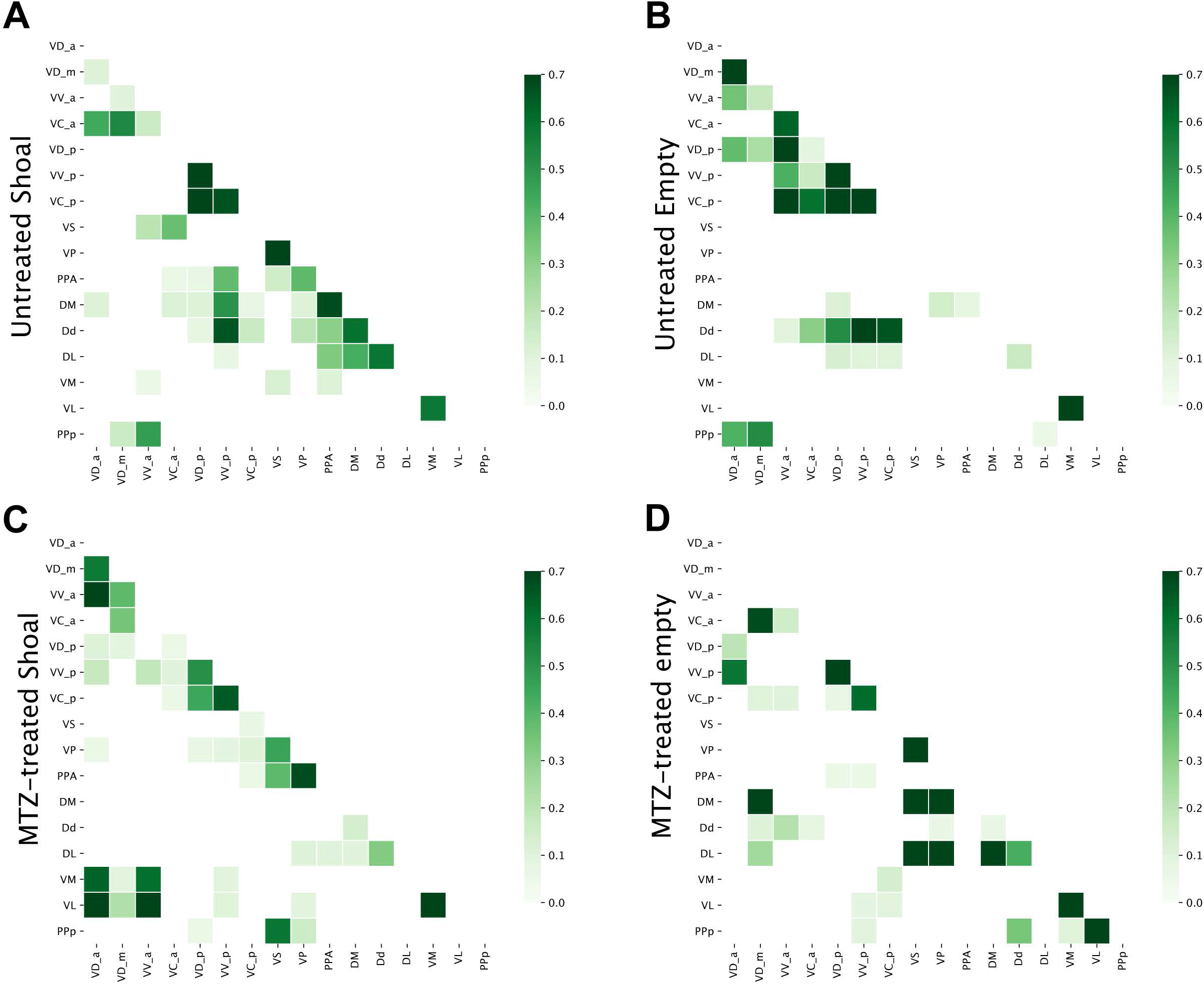
Correlation matrices of functional connectivity for the four treatments reveal early-life OXT shapes social information processing. Regional correlation values are computed joining data from both hemispheres. Resampled correlations were kept if significant (p<0.05). For visualisation purposes, we show only entries of significant correlations higher than 0.05.

**Figure 6-4.**
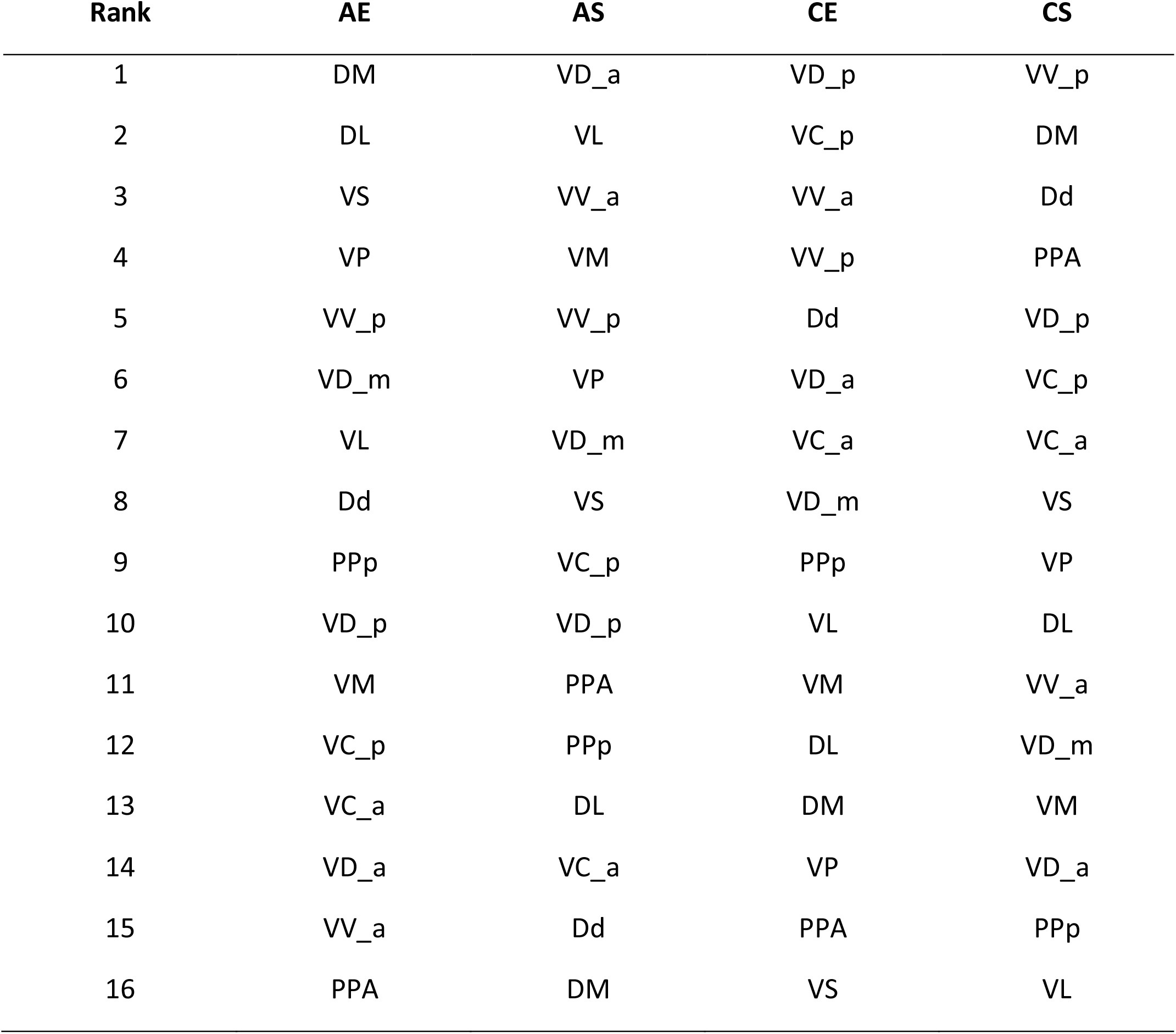
Rank of node strength centralities of adult treated Vs untreated zebrafish, exposed to a conspecific shoal or to empty compartments. *AE*, early-MTZ treated fish exposed to empty compartment; *AS*, early MTZ-treated fish exposed to shoal; CE, untreated fish exposed to empty compartment; CS, untreated fish exposed to shoal; *Vv_a*, anterior part of the ventral nucleus of the ventral telencephalic area (V); *Vv_p*, posterior part of the ventral nucleus of the ventral telencephalic area (V); *Vd_a*, anterior part of the dorsal nucleus of V; *Vd_m*, medial part of the dorsal nucleus of V; *Vd_p*, posterior part of the dorsal nucleus of V; *Vc_a*, anterior part of the central nucleus of V; *Vc_p*, posterior part of the central nucleus of V; *Dm*, medial zone of the dorsal telencephalic area (D); *Dl*, lateral zone of the dorsal telencephalic area (D); *Dd*, dorsal zone of the dorsal telencephalic area (D); *Vs*, supracommissural nucleus of V; *Vp*, postcommissural nucleus of V; *Ppa*, anterior part of the parvocellular preoptic nucleus; *Ppp*, posterior part of the parvocellular preoptic nucleus; *VM*, ventromedial thalamic nucleus; *VL*, ventrolateral thalamic nucleus;

